# Non-saturating dimensionality, contextual dependence, and the limits of unsupervised decoding in motor cortex

**DOI:** 10.64898/2026.01.26.701668

**Authors:** Michael P Silvernagel, Alice Y Tor, Elizabeth J Jun, Stephen E Clarke, Robert Sutherland, Kenji Marshall, Yuxin Wu, Muhammad U Abdulla, Nir Even-Chen, Paul Nuyujukian, Brain Interfacing Laboratory

## Abstract

Understanding how motor cortex generates movement is a foundational challenge in neuroscience. Unsupervised dimensionality reduction techniques, such as principal component analysis (PCA), are widely used to transform high-dimensional neural recordings into a compact, low-dimensional space. The dimensionality of this space—that is, the number of principal components needed to explain a fixed fraction of variance—is broadly assumed to be an intrinsic property of the underlying neural dynamics, potentially modulated by task complexity. Here, by comparing constrained reaching and unconstrained naturalistic behaviors recorded from the same animal on the same day, we show that this assumption breaks down in two distinct ways. First, across four non-human primates, the dominant axes of low-dimensional neural activity separate behavioral contexts rather than movement kinematics, with neural activity shifting rapidly between task-specific regions of state space at task transitions. Notably, traditional dimensionality metrics are insensitive to movement complexity across tasks. Instead, unsupervised dimensionality scales with the number of recorded neurons, exhibiting non-saturating growth up to 1000 simultaneously recorded electrodes, a pattern that holds across PCA, factor analysis, shared variance component analysis, and nonlinear autoencoders. This scaling has direct consequences for decoding: while decoders trained on unsupervised subspaces improve only modestly with electrode count, supervised methods leverage additional electrodes to separate neural states from a vanishingly small fraction of total variance (<10% at 1000 electrodes). Together, these results challenge current views on cortical dimensionality, reveal a greater-than-appreciated role for behavioral context in shaping motor cortical activity, and motivate careful consideration of computational methods as experimental data volumes scale.

## Introduction

The quest to understand how the brain generates movement has a rich history, dating back over a century [18, 19, 30, 59]. For decades, this scientific endeavor framed individual neurons as encoders for specific movement parameters. Landmark studies in primates correlated firing rates of individual motor cortex neurons with kinetic variables like force [7, 17, 61, 64] and kinematic variables such as position, velocity, and direction of hand movement [27, 28, 40, 44]. However, limitations with this approach became apparent; the tuning of single neurons proved bewilderingly complex and context-dependent, failing to provide a unified principle of motor cortical function [8, 57].

This led to studies proposing an alternative hypothesis: motor cortex is not a collection of independent encoders, but a recurrently connected dynamical system that generates motor commands through the collective activity of the entire neural population [10, 11, 58, 66]. In this modern view, the coordinated activity of millions of neurons is constrained to a much lower-dimensional subspace, or “neural manifold” [21, 51]. Understanding the intrinsic dimensionality of the system—the true number of degrees of freedom required to describe the population activity—thus became a central goal [14, 31].

Quantifying dimensionality is critical because it reflects the complexity of the neural computation; a low-dimensional system implies that the brain simplifies the command problem by operating on a compact set of fundamental activity patterns rather than controlling each neuron independently [24]. Unsupervised dimensionality reduction techniques have long been used to simplify analysis of high dimensional neural data [14]. Applying these methods, previous primate studies calculated the fewest dimensions needed to explain a desired variance or performance threshold, estimating the dimensionality of primary motor cortex (M1) to be between 5 and 12 dimensions [10, 22, 46, 70]. Building on these findings, studies have assumed dimensionalities ranging from 8 - 20 to inform methods like jPCA, demixed PCA, factor analysis, latent-state linear dynamical systems, and canonical correlation analysis, all of which require dimensionality to be specified *a priori* [11, 23, 33, 35, 51]. Non-linear models have also emerged as methods to find low-dimensional latent embeddings in neural data [29, 43, 55], but these models typically have not been used to estimate dimensionality in motor cortex.

These studies have supported arguments that motor cortex and other brain regions can be well-represented by low-dimensional neural manifolds [36, 47, 52]. It is important to note, however, that analyses have historically restricted movement to minimize confounding variables [54, 56] and have utilized technologies that provide on the order of 100 electrodes [42]. Theoretical work has proposed that the dimensionality observed in neural recordings may be limited by the complexity of the task itself [25, 62]. This hypothesis suggests that simple, stereotyped laboratory tasks may artificially constrain our view of motor cortical activity, and that the true computational capacity of the system will only be revealed by more complex, naturalistic behaviors [12, 47]. Here, we leveraged advances in wireless neural recording to test this hypothesis. Using two non-human primate cohorts with 96 and 1024 electrodes of recording, we compared same-day recordings of seated reaching tasks and periods of unconstrained, naturalistic activity. With this data, we set out to directly test how the intrinsic dimensionality of motor cortical dynamics scales with movement complexity.

## Results

### Behavioral context generates low-dimensional state space offsets

We investigated open questions regarding motor cortical dimensionality using two non-human primate cohorts. The first cohort consisted of two adult male rhesus macaques (O and U, *Macaca mulatta*), each with 96-channel electrode arrays (Blackrock Microsystems, Salt Lake City, UT) implanted in motor regions of cortex. On the same day, each animal performed two tasks requiring different levels of movement complexity. While seated with its head, legs, and non-reaching arm gently constrained, the animal first performed center-out reaches to one of eight radially-spaced targets (Figure 1A, see Methods). Both reaching periods and periods where all limbs were at rest were noted. Approximately 30 - 45 minutes following the completion of the reaching task, the animal was placed in a large, specialized enclosure for unconstrained behavioral monitoring [60]. Here, the animal traversed the space for treats, generating stereotyped periods of walking, reaching, turning, and sitting (Figure 1B).

**Fig. 1:**
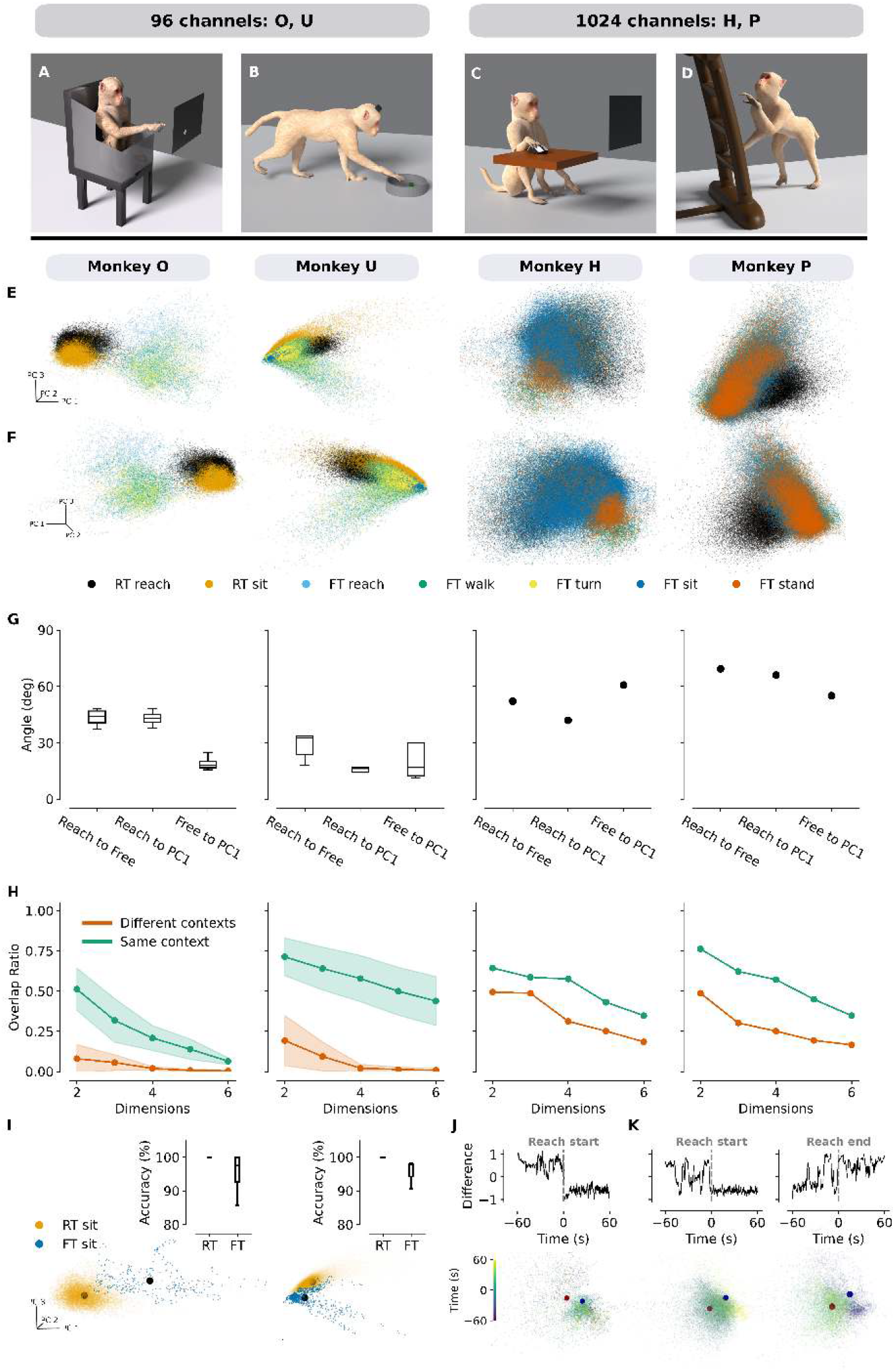
Low-dimensional state comparisons across tasks of increasing complexity. (**A**) Head-fixed radial-8 reaching task (96-channel Utah array). Starting at the center of the screen, the animal reaches for one of eight equally spaced targets. Movement is restricted to the limb of interest to minimize confounding variables. (**B**) Walking and reaching task (96-channel Utah array). Without physical constraints, the animal walks to bowls on opposite sides of a 4.5-m^2^ enclosure to reach for treats. (**C**) Seated computer mouse task (1024-channel Neuralink N1). In a seated position without physical constraints, the animal uses a computer mouse to reach for targets appearing randomly on a 35×35 grid. (**D**) Foraging task (1024-channel Neuralink N1). The animal explores its home environment and forages for treats in an unstructured fashion. (**E**) PCA projection of tasks. For Monkeys O and U, radial-8 and walking-and-reaching data are combined for the projection (O240718, U240614). These data include reaching and sitting (radial-8) and reaching, walking, turning, and sitting (walking-and-reaching). For Monkeys H and P, computer and foraging tasks are combined (H240729, P240608), including reaching (computer task) and walking, sitting, and standing (foraging). Activity legend appears below plots (RT: reaching task; FT: free task). (**F**) Same as (**E**), but rotated 180^°^ with respect to PC 3. (**G**) Angles between main axes of variance. Principal 3-dimensional axes of variance were determined for the reaching and free tasks, and the angles between these axes and PC 1 are shown. These axes do not align with PC 1, contrasting with findings in [35] describing PC 1 as a condition-invariant component tied to movement onset. (**H**) Convex hull overlap. Convex hulls were constructed to encapsulate state space activity for a particular behavior. For Monkeys O and U, these behaviors include reaching activity across different contexts (radial-8 vs. walking-and-reaching task) and reaching/walking behavior within the same context (walking-and-reaching task). For Monkeys H and P, these behaviors include seated reaches vs. unconstrained sitting (different contexts) and sitting vs. walking (same context). Shading indicates 95% confidence intervals. Higher overlap for same-context activities indicates these behaviors are closer in state space than movements across contexts. (**I**) Seated comparison across tasks. Colored dots are the same as in (E) but show only seated periods without limb movement in the head-fixed and unconstrained setups; black dots show centroids of activity. While these behaviors are similar, the specific task context introduces a noticeable state-space offset. Inset shows LDA decoding accuracy on individual time points, highlighting their separability. (**J**) Activity spanning start and completion of the computer mouse task. Red dots represent the centroid of low-dimensional neural activity during the computer task; blue dots mark the centroid for the unconstrained foraging task. Difference in centroid distance is shown above, where positive values indicate activity closer to the foraging centroid and negative values mark activity closer to the mouse task centroid. In monkey H, activity shifts toward the computer task centroid at task onset. (**K**) Same as (**J**) for monkey P. Upon task completion, activity shifts back toward the foraging centroid.

The second cohort, also two adult male rhesus macaques (H and P, *Macaca mulatta*), were implanted with a wireless 1024-channel system (N1, Neuralink, Fremont, CA) in arm-related areas of motor and premotor cortex [41]. These animals performed tasks similar to those studied in Monkeys O and U, each requiring various levels of movement complexity. First, Monkeys H and P performed a seated, mouse-controlled grid task (Figure 1C). Here, each animal used its right arm to move a computer mouse in order to select highlighted squares on a 35×35 grid displayed on a nearby laptop screen. Approximately thirty minutes after the conclusion of that task, the animal performed an unconstrained foraging task, where the animal engaged in various behaviors—including standing, sitting, and walking—as it searched its home enclosure for treats (Figure 1D). A breakdown of behaviors observed across recording sessions can be found in Table S1, and firing rate analysis across sessions can be found in Figure S1.

For each animal, we combined equal amounts of data from both tasks and projected this data into a shared, low-dimensional space using principal component analysis (PCA) (see Methods). Example data from a single day of recording are plotted in the top 3 PCs for each animal in Figures 1E-F. The greatest driver of modulation in this low-dimensional space is not movement complexity, but rather behavioral context, which we broadly define as the collective set of environmental stimuli and motor outputs associated with a given experimental task.

We quantified this modulation by taking the projected neural data for the seated-reaching and free movement tasks, and for each task, finding the main axis of variance in the low-dimensional space. The angle of separation between these axes is plotted in Figure 1G. To avoid the near-orthogonality that arises generically among vectors in high-dimensional spaces, we restricted our angle calculations to the top 3 PCs [13]. For Monkeys O and U, we calculated these values over 4 recording sessions; for Monkeys H and P, these values were calculated for a single session. Across all 4 animals, these axes were not aligned, with median values of 44° (33°) for Monkey O (U) and values of 52° (69°) for Monkey H (P). Interestingly, none of these axes of variance were directly aligned with PC 1, with median angles ranging from 16° to 43° for Monkeys O and U, and angles ranging from 42° to 66° for Monkeys H and P. This stands in contrast to previous work, which proposed that PC 1 is a condition-invariant signal tied to movement onset [35]. Under their hypothesis, the dominant axis of variance for any given task should align with this condition-invariant dimension, a prediction that, based on our findings, does not generalize to unconstrained behavior.

To further quantify this low-dimensional context separation, we explored how kinematically similar movements across contexts compared to more dissimilar movements within a context. For Monkeys O and U, we compared head-fixed reaches to unconstrained reaches for treats (different contexts), as well as unconstrained reaching and walking behavior (same context). For Monkeys H and P, we compared seated computer mouse reaches to periods of unconstrained sitting (different contexts), as well as unconstrained sitting and walking behavior (same context). Sweeping across the top 2 to 6 dimensions, we constructed convex hulls that encapsulated the state space activity for these behaviors and computed the ratio of hull overlap to the total volume occupied by both hulls (Figure 1H, see Methods). Across all 4 animals, the hull overlap was significantly greater for movements within a particular context than for movements across contexts.

Data recorded for Monkeys O and U allowed for more direct movement comparisons across tasks. Across reaching and free movement tasks, periods where the animals were seated with minimal limb movement were compared. Figure 1I highlights these seated periods from Figure 1E. Data were balanced across sessions and projected onto the top 20 PCs, consistent with dimensionalities used in prior studies [34]. Using this data, we constructed a linear discriminant analysis decoder. Decoding was performed on individual 15ms bins of neural activity, with accuracy taken as the median value from a 10-fold cross-validation. Given the high degree of state space separation between contexts, this decoder performed exceedingly well across sessions: for Monkey O (U) the decoder had a median accuracy of 100% (99.9%) classifying the reaching task data and 99.9% (97.8%) classifying free task data.

For Monkeys H and P, we observed shifts in state space activity as the animals moved between contexts. Before and after completing the reaching task, Monkeys H and P returned to the area of their enclosure where the free movement task occurred. After finding the state space centroids for reaching and free task data, we calculated the 20-dimensional distance for 60 second periods encompassing the start and end of the reaching task (see Methods). We then calculated the difference between the reaching and free centroid distances, and the result was smoothed using a one second (66 15ms bins) rolling window. This difference yielded an intuitive metric: positive values indicate activity closer to the free task centroid, while negative values indicate activity closer to the reaching task centroid. Figure 1J illustrates this transition as Monkey H began the reaching task. Upon task onset, the difference metric shifted from positive to negative, indicating that the neural population activity migrated closer to the reaching task centroid. This transition is also evident when examining the temporal progression of neural data within the top three PCs. Figure 1K displays this progression for Monkey P as the animal initiated and concluded the reaching task. Neural activity transitioned toward the reaching task centroid at the start of the session; conversely, once Monkey P finished the task and returned to the main area of the enclosure, the activity reverted toward the free task centroid.

### Neural dimensionality scales with electrode count

We next investigated the effect of movement complexity on neural dimensionality. For each animal, we separately performed PCA on neural activity from the seated reaching and free movement tasks, and then calculated the number of PCs needed to account for 80 percent of the total variance (Figure 2A, see Figure S2 for individual recording sessions). We then repeated this calculation after combining equal amounts of reaching and free movement data. Across all animals, dimensionality did not drastically differ between the reaching task, movement task, and combined datasets. In contrast, we observed a sharp increase in dimensionality when scaling from 96 to 1024 recording channels. For Monkey O(U), dimensionality ranged from 31 to 37 (36 to 38); for Monkey H(P), dimensionality ranged from 186 to 218 (163 to 173).

**Fig. 2:**
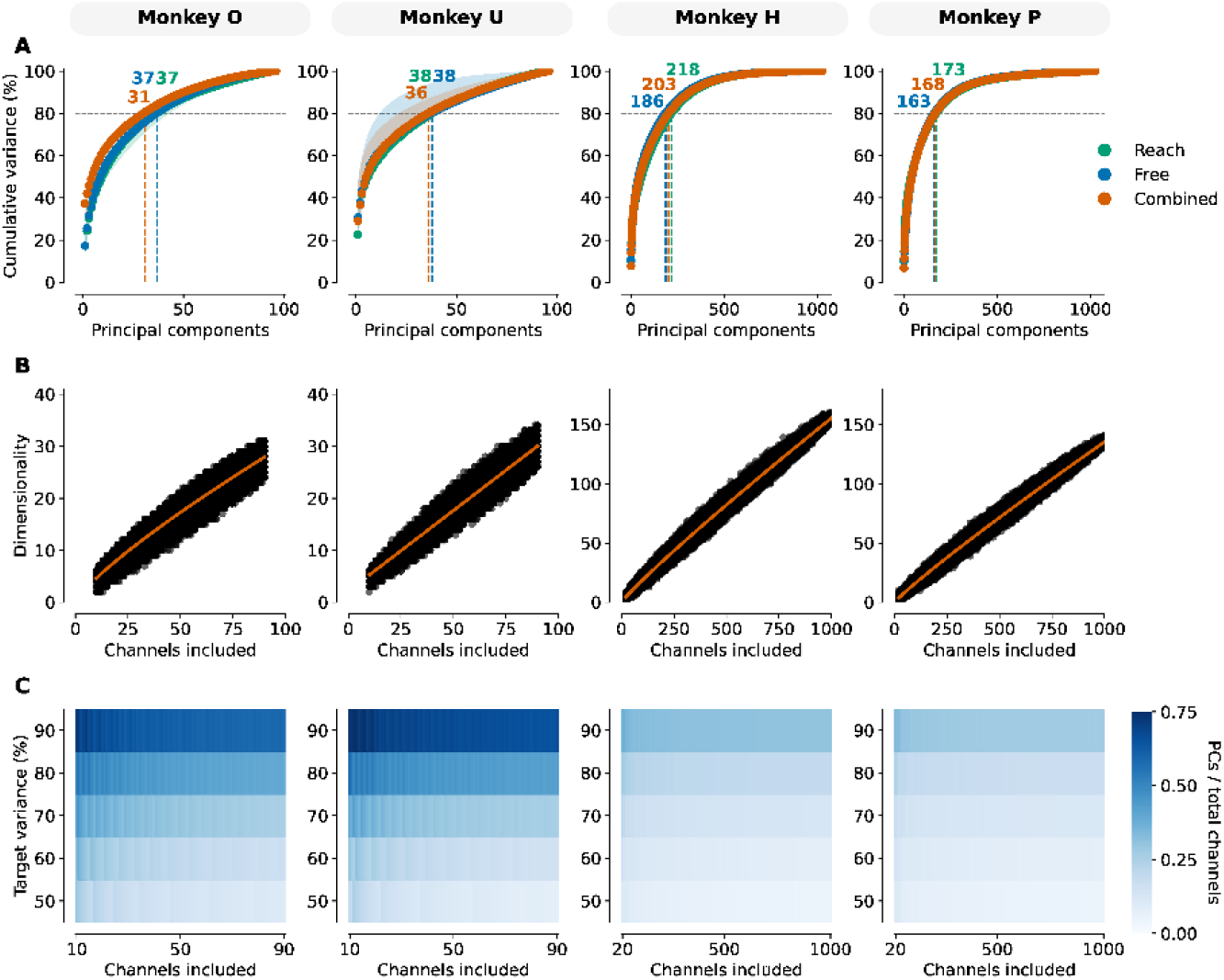
PCA-dimensionality scaling with increasing electrode count. **(A)** Cumulative variance explained plots for reaching task (green), freely-moving task (blue), and combined reaching/freely-moving tasks (orange). Dots indicate median values across sessions, and shading reflects the interquartile range. Vertical dashed lines indicate the number of principal components needed to explain 80 percent of the underlying variance, and the dimensionalities corresponding to these lines are shown at the top of the plot. While the freely-moving tasks require the animals to engage in more complex behavior, the dimensionality between these tasks and the seated reaching tasks is relatively unchanging. However, as electrode count jumps from 96 channels (Monkeys O and U) to 1024 channels (Monkeys H and P), PCA-based dimensionality increases dramatically. **(B)** Dimensionality as a function of channels included in analysis. Across all monkeys, as the number of electrodes increases, dimensionality does not plateau, as would be expected if dimensionality were an intrinsic property of the data. Rather, dimensionality grows as the analysis scales to 1000 channels. Fit lines represent the linear or power law model that best described the data (see Table S2). If neither model performed significantly better than the other, a linear fit is used. **(C)** Heatmaps showing principal components needed to explain a fixed percentage of variance as a ratio of electrodes included. Across different variance thresholds, the ratio of PCs to the channels included in the analysis holds relatively constant. This is consistent with the scaling relationship shown in (B).

To explore the relationship between channel count and dimensionality, we repeated the dimensionality calculations using random subsets of electrodes. For Monkeys O and U, we sampled electrode counts ranging from 10 to 90; for Monkeys H and P, we sampled from 10 to 1000 in increments of 10. Each subset size was resampled 1000 times with replacement, and the resulting dimensionality estimates were regressed to determine a line of best fit. Figure 2B shows these estimates on the free movement task; seated mouse task estimates can be found in Figure S3A. For most animals and tasks, power law fits (D ∝ N^*λ*^), with *λ* valued at 0.8 for Monkeys O and U and 0.9 for Monkeys H and P) significantly outperformed linear fits (two-sided Vuong’s closeness test, see Tables S2, S3). However, power law and linear models generated nearly identical *R*^2^ values, necessitating greater electrode counts to draw definitive conclusions.

To verify our dimensionality result, we repeated this analysis using three additional methods. Since increasing electrode counts can introduce independent noise sources that may inflate PCA-based dimensionality estimates, we calculated dimensionality using factor analysis (FA) and shared variance component analysis (SVCA), two unsupervised methods that isolate shared from independent variance. Like PCA, however, FA and SVCA are linear methods, which may overestimate dimensionality when the true underlying architecture is nonlinear [2, 15]. We therefore also estimated dimensionality using autoencoder models, which offer a nonlinear analog to PCA (see Methods). Across all three methods, we observed scaling relationships similar to those uncovered by PCA, suggesting that the observed scaling is driven by structured, shared variance rather than independent noise (Figure S4). The autoencoder estimates exhibited substantially greater variance than the linear methods, yielding weaker linear fits—particularly for Monkey O(U), where the scaling relationship was largely absent (*R*^2^ = 0.12(0.11))—likely reflecting sensitivity to initialization and local optima in the low-channel regime. Monkey H(P) retained moderate fits (*R*^2^ = 0.50(0.58)), though these were notably weaker than those obtained with FA and SVCA. Despite this added variance, the autoencoder estimates did not reveal a qualitatively different scaling behavior, consistent with the interpretation that the dimensionality scaling we observe reflects genuine structure in the shared neural variance.

We further verified that this effect was not an artifact of a particular target variance. Figure 2C repeats the previous channel sweeps across different variance thresholds (see Figure S3B for seated mouse task). Here, heatmap values reflect the median ratio (number of PCs /subset size) needed to achieve that variance threshold. For all animals across all thresholds, these ratios are nearly constant, mirroring the relationship found in Figure 2B.

When analyzing general behavior on the timescale of minutes, PCA-derived dimensionality estimates do not readily distinguish between movements of varying complexity. However, isolated movements drawn from the same behavioral context may be differentiated by dimensionality. For all animals, we manually identified distinct movements during the period of free activity. For Monkeys O and U, these activities included periods of sitting, walking, reaching, and turning during an unconstrained walking and reaching task; for Monkeys H and P, we marked periods of sitting, standing, and walking during unconstrained foraging (see S1, Methods). For each recording session, neural data corresponding to these movements were partitioned into 500ms epochs; movements with fewer than 60 epochs were excluded from that session’s analysis. These epochs were randomly sampled and concatenated to generate a 30 second period on which we calculated PCA dimensionality. We repeated this process 1000 times for each behavior, creating dimensionality distributions for each movement (Figure S5).

For comparison, we calculated the inverse compression ratio (ICR, see Methods)—a metric which has been deployed for seizure detection—on raw spiking data from the same 30 second windows used for PCA calculations [69]. These values are shown in Figure S6. As resamples drawn from the same session are not statistically independent, we used Cliff’s delta as a non-parametric effect size for each within-session pairwise comparison, reporting magnitude thresholds (small, medium, large) as defined in [49]. Within a given session, both ICR and PCA dimensionality typically distinguished movement types with medium-to-large effect sizes. For Monkeys O and U, ICR yielded fewer negligible effects across sessions, with only walk vs. turn events on two Monkey U sessions returning a negligible effect. Although ICR and PCA may distinguish movement types within a given recording context, neither generalized across animals or recording modalities. Absolute ICR and PCA values varied across animals and sessions, and the rank ordering of movements was not preserved across animals; for example, relative to other movements, both ICR and PCA produced lower values for sitting in Monkey U, but higher sitting values for Monkeys H and P. What meaning, if any, can be ascribed to a particular ICR or PCA value remains a subject for future research.

### Supervised learning improves decoding performance using a small fraction of data variance

With the advent higher electrode count recording modalities, the degree to which these devices can be leveraged to improve movement decoding remains unanswered. To probe this question, we examined the seated reaching task performed by all animals. Monkeys O and U performed a widely studied radial-8 reaching task [9, 10]. Here, the animals were gently constrained except for the arm contralateral to the implant location, and they performed repeated, stereotyped reaches to eight equally spaced targets. Monkeys H and P were also seated but had no physical constraints. Using a cursor controlled by a computer mouse, they selected random targets on a 35 × 35 square grid (see Methods). To compare this task with the radial-8 reaches performed by Monkeys O and U, we defined eight equally spaced acceptance windows, each spanning 40° and centered on the standard radial-8 target locations. Cursor movements falling within these windows were pooled, and reaches were decoded by direction. While this pooling facilitated direct comparison, the grid task presented a more challenging decoding problem because, unlike the stereotyped radial-8 reaches, grid reaches within a class varied in angle, length, and duration.

To determine the impact of channel count on PCA decoding performance, we trained a Gaussian Naive Bayes decoder on neural trajectories composed of the top 20 PC dimensions. This choice of dimensionality aligned with previous literature and represented the threshold above which decoding performance decreased (Figure S7) [33]. Decoded trajectories began 60 ms after reach onset and extended up to 510 ms for Monkeys O and U or 330 ms for Monkeys H and P. We swept both the length of the decoded trajectory and the number of channels used, computing decoding performance across 1000 random channel subsets for each trajectory-length and channel-count pair (see Methods). Figure 3A shows mean decoding performance across channel samples and recording sessions (individual sessions for Monkeys O and U are shown in Figure S8A). Across both tasks, PCA decoding performance improved as trajectory length and channel count increased, with peak performance reaching 58% (79%) for Monkey O (U) and 75% (73%) for Monkey H (P).

**Fig. 3:**
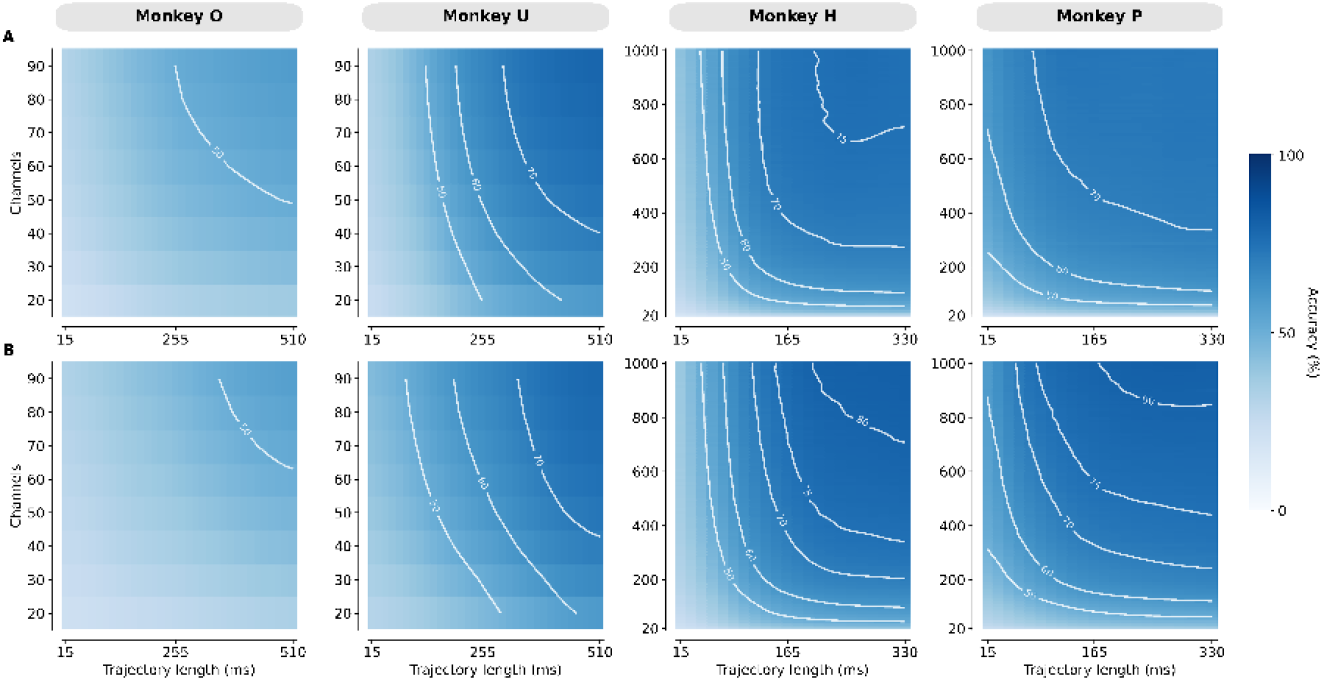
PCA and mcLDA decoding for reaching tasks. **(A)** Heatmap of PCA decoding performance. For all animals, increasing electrode count and trajectory lengths result in modest performance increases. **(B)** Heatmap of mcLDA decoding performance. For Monkeys O and C, mcLDA decoding performance is comparable to PCA. For Monkeys H and P, mcLDA outperforms PCA at higher channel counts. The improvement over PCA is most pronounced for trajectory lengths above 165 ms and channel counts above 400.

We repeated this decoding analysis using multi-class linear discriminant analysis (mcLDA), a supervised method that finds a low-dimensional projection by maximizing between-class variance while minimizing within-class variance (see Methods). With eight reach directions, mcLDA yielded a 7-dimensional projection space for our neural data. As with PCA, mcLDA performance improved with increasing integration lengths and channel counts. For Monkey O (U), mcLDA performance was comparable to PCA, with accuracy peaking at 56% (78%)(Figure 3B, see Figure S8B for individual recording days). However, as more channels were added to the analysis, mcLDA outperformed PCA. For Monkey H (P), this performance is particularly noticeable for trajectory lengths above 165 ms and channel counts above 400, with decoding performance peaking at 82% (81%). Notably, mcLDA’s improved decoding relied on a subspace capturing far less of the total variance than PCA. At the maximum available channel counts, PCA explained 68% (62%) of the variance for Monkey O (U) and 42% (44%) for Monkey H (P) (Figure S9A). The corresponding mcLDA values were 9% (12%) for Monkey O (U) and 7% (8%) for Monkey H (P) (Figure S9B). This dissociation between variance and decoding accuracy highlights that the most task-relevant subspace is not always the highest-variance one.

### Increased electrode count improves low-dimensional separability for supervised learning

To investigate how electrode count influences low-dimensional PCA and mcLDA representations, we plotted two-dimensional neural trajectories for the computer mouse task, colored by reach direction. For Monkeys O and U, PCA trajectories (both individual and averaged) using all 96 channels were not separable by direction (Figure S10). After applying mcLDA, individual trajectories remained overlapping, but averaged trajectories became separable by direction (Figure S11).

Similar patterns emerged for Monkeys H and P when constraining the analysis to 100 channels. Increasing the channel count to 1000 revealed that individual PCA trajectories remained inseparable (Figure 4A). While averaging trajectories by reach direction allowed for discrimination above 500 channels (Figure 4B), these trajectories still exhibited significant overlap.

**Fig. 4:**
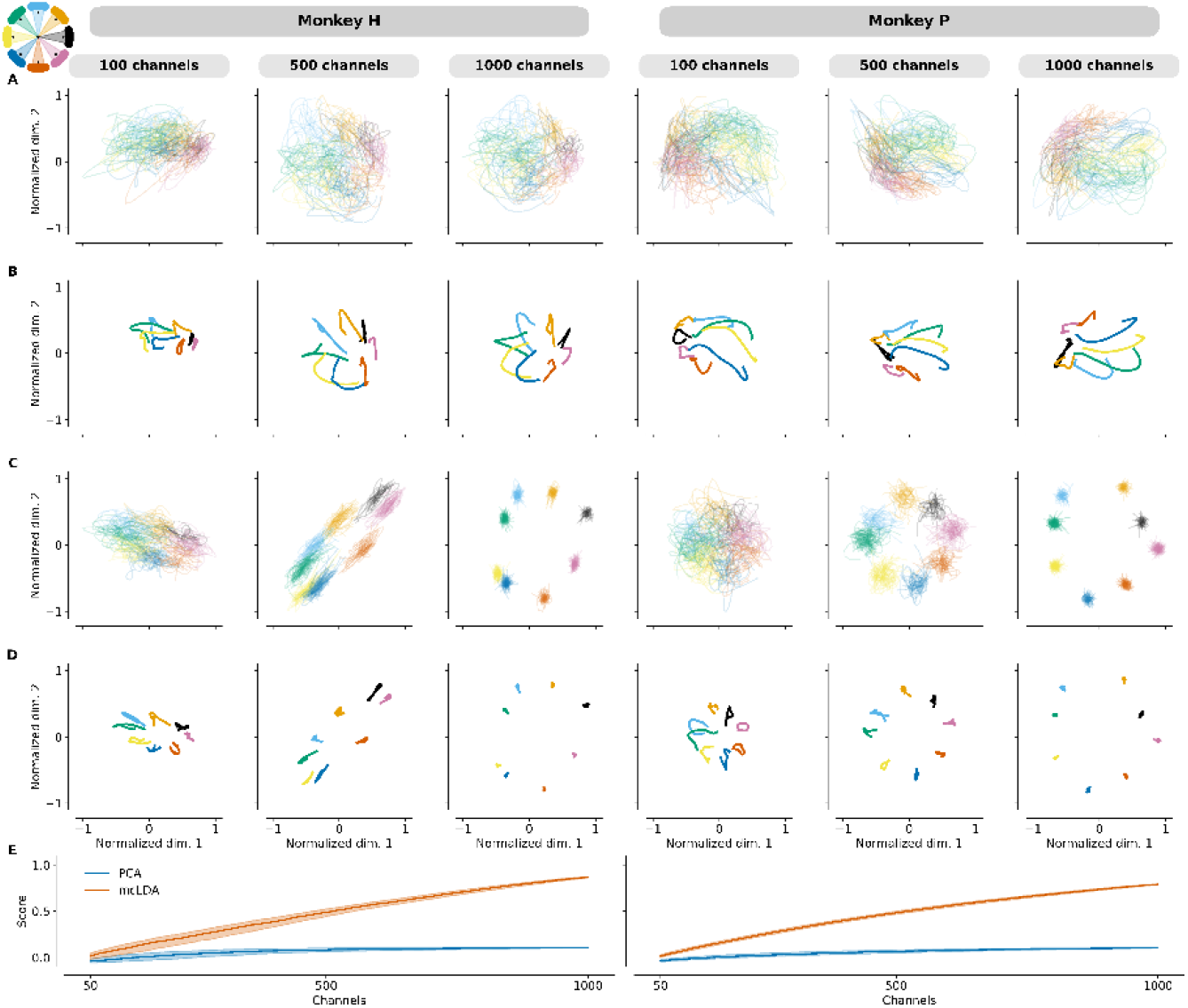
PCA and mcLDA reach trajectories for computer mouse task. Movement angles were calculated for all reaches, and reaches falling within 8 equally spaced 25° windows were grouped together. Trajectory colors correspond to these windows, as shown in the graphic in the upper left corner of the figure. **(A)** Individual PCA reach trajectories. At each channel count, trajectories are generated from a random subsample of electrodes. Even at 1000 electrodes, individual traces cannot be readily separated by movement direction. **(B)** Averaged PCA reach trajectories. These averages are generated from the trajectories shown in (A). As electrode count increases, structure differentiating reach direction appears. However, these averaged trajectories are not fully separable as trajectories still overlap. **(C)** Individual mcLDA trajectories. Trajectories are generated from the same set of randomly subsampled electrodes used in (A). At high channel counts, individual mcLDA trajectories cluster according to reach direction and are highly separable. **(D)** Averaged mcLDA trajectories. Averages are generated from the trajectories shown in (C). These trajectories further highlight the separability achieved by mcLDA, giving insight into how this approach provides superior decoding performance. **(E)** PCA and mcLDA silhouette scores for increasing channel counts. Shading indicates the interquartile range. As channel count increases, mcLDA leverages the additional information to better separate clusters, as evidenced by the increasing silhouette score. For PCA, this score improves modestly, plateauing at a score of 0.1 for both animals.

In contrast, mcLDA yielded clear clusters at 500 channels even for individual trajectories (Figure 4C). Cluster separation continued to improve with higher channel counts and was further enhanced when trajectories were averaged (Figure 4D). To quantify these observations, we calculated silhouette scores for individual PCA and mcLDA trajectories (Figure 4E, see Methods). These scores, ranging from −1 to 1, measure cluster separability by comparing the mean intra-cluster distance to the mean distance to the nearest neighboring cluster. A score approaching 1 indicates distinct, well-separated clusters, while a score near 0 suggests significant overlap.

Trajectories were generated by randomly subsampling electrodes at intervals of 10 channels. This process was repeated for 1000 iterations, and median values are reported. In both animals, PCA silhouette scores improved modestly with increasing channel count before plateauing at approximately 0.1. Conversely, mcLDA scores continued to improve as channel counts increased. This indicates that even for relatively simple movements, increasing electrode density provides additional information that can be leveraged for robust decoding.

## Discussion

Our results challenge a foundational concept in motor neuroscience: that unsupervised techniques can distill the complex activity of motor cortex into an interpretable, low-dimensional manifold. We demonstrate that neural dimensionality, as measured by the canonical method of principal component analysis, is not an intrinsic, fixed property of motor cortex. Instead, it is fundamentally constrained by the scale of observation, increasing with the number of recorded neurons. This aligns with recent studies using calcium imaging in mouse cortex and fMRI in human visual cortex, which noted similar scaling relationships [26, 37, 63]. Consequently, the low dimensionality reported in previous studies may be an artifact of historical technological limitations—specifically, the restricted number of simultaneously recorded electrodes.

The low dimensionality of motor cortex reported in previous work is consistent with the scaling relationship we identified. While the dimensionalities observed in our 96-channel center-out reaching data were slightly higher than those in prior literature, this discrepancy likely stems from differences in thresholding or preprocessing choices. For instance, adopting a 60% variance threshold — consistent with [22] — reduces the reaching dimensionalities for Monkeys O and U to 16 and 13, respectively, aligning closely with prior results (Figure S12). Trial-averaging similarly reduces dimensionality. While trial-averaged dimensionality does scale with channel count, at 96 channels we observe dimensionalities of 3 to 5, extending to 6 to 8 at 1000 channels (Figure S13). Here, trial-averaging acts as a supervised form of dimensionality reduction, with dimensionality limited by task complexity. This is consistent with what we observed with mcLDA (which, for an 8-target reach, has a maximum dimensionality of 7) and aligns with previous work [1].

At lower electrode counts, the number of components needed to explain a significant portion of the variance is small enough to suggest a compact, low-dimensional structure. However, as more neurons are recorded, this simplicity gives way to ever-increasing complexity. This is reflected in the changing structure of the eigenspectrum, where the relative variance captured by the top principal components progressively decreases as channel counts increase (Figure 2A). The near-linear scaling of dimensionality with the number of recorded neurons indicates that, at least up to approximately 1000 electrodes, each newly sampled neuron provides novel, weakly correlated information rather than redundant signals from a small, fixed set of underlying latent variables. Additionally, comparable scaling relationships were observed using FA, SVCA, and autoencoder models, suggesting that this growth in dimensionality is driven by structured, shared variance rather than an artifact of independent noise introduced by additional electrodes. This motivates a re-evaluation of the structure of motor cortex and the information it encodes.

While our nonlinear analyses were restricted to autoencoders, motor cortical dynamics are complex and might be better modeled using a different nonlinear manifold learning or nonlinear dimensionality reduction technique [29, 43, 55]. However, applying these models to large-scale neural recordings introduces several computational and mathematical considerations. Many nonlinear systems rely on distance- or similarity-based reasoning (e.g., *k*-nearest neighbors or radial basis functions). In highdimensional spaces, the mathematical distance between data points becomes highly uniform, causing these similarity metrics to degrade and lose their predictive power. Furthermore, the number of data points required to accurately map these complex nonlinear relationships grows exponentially as the underlying dimensionality increases; for instance, the correlation dimension method—a foundation for many nonlinear algorithms—demands approximately 10^*d/*2^ samples, where *d* is the intrinsic dimensionality [6]. Indeed, a systematic evaluation of dimensionality estimation algorithms on synthetic motor cortical data found that nonlinear methods began to severely underestimate true dimensionality for intrinsic dimensions above approximately 10, regardless of whether the underlying embedding was linear or nonlinear [2]. If the true intrinsic dimensionality of motor cortex is indeed high, as our scaling results suggest, accurately estimating dimensionality using nonlinear techniques becomes challenging. While nonlinear architectures hold promise for uncovering complex neural embeddings, a careful exploration of their utility and limitations in scaling to thousands of channels is outside the scope of this work and should be a dedicated focus of future research.

While the most appropriate method of representing motor cortex remains an open question, examining the dominant linear dimensions can still generate new insights. Specifically, shifts in behavioral context were readily visible in the top principal components of the neural data, a finding that runs contrary to the expectation that dominant patterns of variance are driven primarily by movement kinematics. The influence of context on the low-dimensional state, even for kinematically similar actions, further reinforces the fact that motor cortex activity is not purely motor. There are a number of potential factors driving these contextual shifts, including posture, sensory stimuli, and emotional state. While further investigation is needed to parse out the forces driving these context shifts, our results may point to a greater-than-appreciated integration of sensory and other contextual signals into the generation of motor commands.

The dimensionality scaling we observed also has implications for decoding movement, particularly for brain-machine interfaces (BMIs). If there is no stable, low-dimensional subspace that captures the essence of motor commands, then unsupervised methods like PCA are fundamentally limited. Indeed, our results show that the dimensions critical for decoding movement direction account for a small fraction of the total neural variance as electrode counts increase. This explains why PCA-based decoding improves only modestly with more channels—it is an approach optimized to find the largest signals in the data, but the signals most relevant for decoding task direction are not the largest. Yet this same high-dimensional structure that confounds unsupervised methods is precisely what supervised learning exploits. By leveraging knowledge of the intended movement, supervised methods like mcLDA can identify precise, information-rich subspaces, even if they contribute little to the total measured neural variance. The improvement in decoding performance and state-space separability with mcLDA, especially at higher electrode counts, highlights the key utility of supervised methods: extracting behaviorally relevant subspaces from complex, high-dimensional neural data.

This is not to say that unsupervised approaches are without value. They remain effective tools for identifying the dominant patterns of neural co-variation and, as shown here, can effectively distinguish broad behavioral contexts. The challenge lies in explaining the real world meaning of high-variance patterns of neuronal activity (i.e., eigenvectors or principal components). A path forward may be to ground these statistical descriptions in the physical architecture of the brain. Future studies could combine large-scale recordings with methods that identify cell types or neural circuitry, such as optogenetic tagging of projection-specific neurons or anatomical reconstruction [3, 16, 48]. This could align dimensions with specific, anatomically defined cell classes or projection pathways. Such an approach could imbue the dimensions found by unsupervised dimensionality reduction techniques with mechanistic meaning, revealing how circuit structure gives rise to function.

A critical question for these future large-scale studies is whether dimensionality continues to scale with the number of recorded neurons. Answering this question necessitates higher electrode count recording technologies. Additionally, given the performance gains and increasing state separability achieved by supervised learning as channel count increases, recording technologies that leverage information from a large population of neurons will be crucial to developing precise and robust decoding algorithms. Attempts to decode more complex behavior will likely only further the need for these high electrode count recording devices.

The success of supervised learning to decode complex movements, however, also hinges on the availability of well-labeled behavioral data. Historically, neuroscience has faced a trade-off between studying simple, highly constrained tasks that are easy to label (e.g., center-out reaching) and studying complex, naturalistic behaviors that are challenging to quantify or systematically repeat. This trade-off has limited our ability to understand how the brain generates the rich motor repertoire of daily life. However, recent technologies may help bridge this gap. Markerless motion tracking methods using RGB [4, 38] or depth camera video[60] can give researchers access to full-body kinematics as animals engage in naturalistic behaviors. Using methods like motion sequencing, behavioral phonemes also can be extracted from this video data, providing additional contextual information [5, 32, 67, 68]. By recording from thousands of neurons while animals engage in unconstrained, naturalistic behaviors, and simultaneously capturing detailed kinematic data, we can generate the rich datasets needed to train supervised learning algorithms.

Ultimately, our work issues three calls to action: (1) developing higher electrode count recording devices; (2) grounding unsupervised learning analyses in neural architecture; and (3) pairing supervised learning methods with rich kinematic and behavioral data. By leveraging the unique capabilities of both supervised and unsupervised learning, we believe systems neuroscience can generate new insights into how the brain performs its operations, including the control of movement.

## Acknowledgments

In addition to the authors affiliated with Stanford, members of the Brain Interfacing Laboratory include Michele Wechsler, Alexandra Paraskevopoulou, Mackenzie Risch, Stephen I Ryu, Alissa Ling, Iliana Bray, and Sydney Hunt. We thank Kimberly Chin for adminstrative support.

## Author contributions

MPS designed study framework, led data collection in Monkeys O and U, programmed data processing pipelines, analyzed data, prepared figures, and wrote original paper (conceptualization, investigation, data curation, formal analysis, methodology, software, visualization, writing – original draft). AYT wrote the imaging software used for acquiring data from Monkeys H & P (software), processed the resulting imaging data (data curation), and prepared Figure 1B-D (visualization). EJJ assisted in acquiring data from Monkeys O & U (investigation) and prepared Figure 1A (visualization). SEC assisted in acquiring data from Monkeys O & U (investigation). RS and NEC assisted in data acquisition from Monkeys H & P (investigation). KM wrote data analysis software for Monkeys H & P (software). YW assisted with imaging software for Monkeys H & P (software). MA authored the multiclass LDA software library and code to time-align center-out reaches (software). NEC oversaw research operations with Monkeys H & P (project administration, resources, supervision). PN acquired data from Monkeys H & P and contributed to all aspects of the research (conceptualization, data curation, funding acquisition, investigation, methodology, project administration, resources, supervision, validation). All authors reviewed and edited the paper (writing – review & editing). All members of the Brain Interfacing Lab also provided insight on review and editing.

## Data and materials availability

Data will be made publicly available at the Stanford Digital Repository, with deposition currently pending.

## Methods

### Animal Procedures

All animal procedures and protocols at Stanford were reviewed and approved by the Stanford University Institutional Animal Care and Use Committee (IACUC). All animal procedures and protocols at Neuralink were reviewed and approved by the Neuralink Institutional Animal Care and Use Committee (IACUC).

### Implantation Details - Utah Array

Two adult male rhesus macaques (O and U, *Macaca mulatta*), were each implanted with 96-channel electrode arrays (Utah array, Blackrock Microsystems, Salt Lake City, UT) in motor regions of cortex. Monkey O (age 19) was implanted with two arrays on July 7, 2023. Monkey U (age 14) was implanted with three arrays on August 4, 2017. All arrays were implanted in arm regions of motor cortex, as determined by visual anatomical landmarks using standard neurosurgical techniques.

### Implantation Details - N1

Two adult male rhesus macaques (P and H, *Macaca mulatta*), were each implanted with 1024-channel wireless recording systems (N1, Neuralink, Fremont, CA). Electrodes were distributed throughout regions of primary motor and premotor cortex. These systems were implanted according to proprietary neurosurgical techniques and procedures developed by Neuralink.

### Head-fixed Reaching Task

Details regarding the head-fixed experimental setup and reaching task have been previously described [9, 10]. In brief, the animal was gently constrained in a seated position with the arm contralateral to the implant location allowed to move freely. A reflective bead was placed on the third and fourth proximal phalanges, and an infrared camera (Polaris Spectra, Northern Digital Inc., Waterloo, Canada) optically tracked three-dimensional movements at 400 Hz. The animal operated in a virtual workspace, where the bead’s position was projected onto a computer screen so that when the animal extended the arm straight out, the cursor mapped to the origin (0,0). This ensured centered visual alignment without having to reach across the body. The task also enforced a maximum permitted distance from the screen along the axis orthogonal to its surface, requiring the monkeys to reach fully outward and make stereotyped movements within a defined work space of approximately 0.25m^3^.

The animal performed a center out reaching task. A trial would begin when a green target (15mm diameter) at the (0,0) coordinate of the task workspace. When the animal moved the cursor to the target and held for the required hold time of 500ms, one of eight equally spaced circles distributed over an invisible circle of radius 100mm would randomly appear. Upon moving and holding the cursor at the new circle, the animal received a juice reward, and a new trial would begin with the center target reappearing. Although target locations were randomized over sequential trials, over the recording session, targets were equally distributed so that a similar number of reaches was performed to each target.

Cursor positions provided by the Polaris camera were first low-pass filtered (8th-order Butterworth, 20 Hz cutoff), and acceleration was approximated by applying a Savitzky-Golay filter (2nd-order, 100 ms window) to the smoothed data. Reach start times were taken as the point of maximum acceleration following target onset. These reaches were performed over sessions ranging from 30 - 60 minutes in duration.

### Unconstrained Walking and Reaching Task

A detailed overview of the unconstrained behavioral setup has been previously reported [60]. This enclosure, measuring roughly 2.1×2.1×2.4 m (l×w×h), was outfitted with 4 Azure Kinect RGB-D cameras (Microsoft, Redmond, WA) that generated high resolution depth and color images at a rate of 30Hz. Neural data was wirelessly transmitted (CerePlex W, Blackrock Microsystems, Salt Lake City, UT) to an array of 16 panel antennas (PA-333810-NF 3.3GHz-3.8GHz 10dBi Panel Directional Outdoor Antenna, FT-RF, Jhubei City, Taiwan) located outside the enclosure. Image data and neural data were synchronized by routing the trigger signal generated by the Kinect cameras to the same acquisition system (Cerebus, Blackrock Microsystems, Salt Lake City, UT) used to process all wireless neural data. This synchronization was validated by a set of three LED lights located outside the enclosure which encoded a binary signal (corresponding to the on/off status of the lights) into the RGB frames. The trigger signal governing light activity was also directed to the neural acquisition system, allowing for direct alignment of the neural activity and RGB frames.

While in the unconstrained rig, the animal engaged in a walking and reaching task. Upon entering the enclosure, an auditory cue would direct the animal to one of two bowls located on opposite ends of the space. The animal would walk to the bowl and reach for a treat, at which point a second auditory cue would alert the animal to the presence of a treat in the other bowl, prompting the animal to turn around and repeat the task. In this way, stereotyped sequences of walking, reaching, and turning behavior were produced, along with periods of sitting when the animal rested. All behaviors were tagged by visual inspection of the RGB video data.

Walking behavior was defined by periods where the animal was facing the target bowl, had its shoulders perpendicular to the target, and was continually advancing forward towards the treat. This continued until the beginning of the reach period, which lasted from the point when the animal lifted its reaching arm off the ground, acquired the treat, and delivered the treat to its mouth. Periods of turning followed treat acquisition and lasted from the time the animal began turning its body until the point its shoulders were perpendicular to the direction of the target bowl on the opposite side of the enclosure. Sitting periods were noted whenever the animal was in a seated position with all four of its limbs at rest.

The walking and reaching task always followed the head-fixed reaching task, with approximately 30 minutes separating these tasks. The walking and reaching tasks lasted approximately 20 - 45 minutes.

### Unconstrained Mouse-Controlled Grid Task

Here, the animal performed a mouse-controlled grid task. While in a seated position, the animal used its right hand to move a computer mouse; apart from the limb used to control the mouse, all other limbs were at rest. This mouse controlled a cursor that displayed on a nearby laptop screen. The cursor was used to select a highlighted square on a 35 × 35 square grid. Successful trials occurred when the animal used the cursor to hover over the highlighted square for a predetermined hold time. Successful trials were accompanied by a liquid smoothie reward. The animal’s movement was tracked by depth cameras (see Improved Depth Camera Setup) and the position of the cursor on the screen. Cursor positions were sampled at 1 kHz. These data were first low-pass filtered (8th-order Butterworth, 20 Hz cutoff), and acceleration was approximated by applying a Savitzky-Golay filter (2nd-order, 100 ms window) to the smoothed data. Movement onset times were taken as the point of peak acceleration following target appearance. Taking the vector formed by the cursor start position and the location of the target, reach angle was then determined as the clockwise angle formed between the vector and the y-axis. Reaches were classified as belonging to a given direction if they fell within one of eight 25° windows. These windows were centered at points starting at 0° and incrementing by 45°, mirroring the location of the radial-8 targets. These reaches were performed over sessions lasting approximately 20 minutes.

### Unconstrained Foraging

Using depth camera images (see Improved Depth Camera Setup), behavior was monitored as the animal foraged for treats in its home enclosure. Enrichment was randomly scattered on the enclosure’s floor, which was also mixed in with hay. Additionally, several objects and toys in the enclosure were smeared with honey. The animal was then allowed to forage for treats and lick honey off objects in an unstructured fashion. Periods of walking, sitting, and standing were manually labeled by visual inspection.

### Improved Depth Camera Setup

For tasks involving animals with the N1 systems, an improved depth camera setup was developed to monitor behavior. Four Helios2+ Time of Flight (ToF) IP67 depth cameras (LUCID Vision Labs, Richmond, Canada) were mounted on tripods at heights between 2 and 6 feet, arranged around the animal’s housing enclosure. Cameras were intentionally mounted at unaligned positions to capture the largest scope and variety of animal movements; specific positions varied between each recording session. The depth cameras collect high-resolution depth images (640 x 480 pixels, 16-bit single-channel pixels) at a frame rate of 90 Hz. Each frame was given a nanosecond-precision timestamp, enabled by Precision Time Protocol (PTP).

Each camera was operated with a customized voltage-gated trigger line. The trigger signal was generated using built-in LED control functions on a wESP32 microcontroller (Silicognition LLC, Longmont, CO) in sequence with a TLV2372 operational amplifier (Texas Instruments, Dallas, TX) in negative feedback to produce a highly temporally consistent 90 Hz 12V square wave. The cameras are configured to drop triggers if they coincide with another frame acquisition (the depth cameras acquire at each trigger; the RGB cameras acquire every 6 triggers). Because each depth camera relies on emitting and recording infrared radiation, each camera is offset from the last by 1 ms, so for every 10 ms of acquisition time, 4 depth images were acquired. There was no temporal offset applied to RGB cameras.

All cameras received power over Ethernet (PoE) and streamed their data via Ethernet through a MikroTik netPower 16P Ethernet switch (MikroTik, Riga, Latvia) to a connected desktop machine. This desktop machine (Hewlett-Packard, Palo Alto, CA) ran Ubuntu 22.04 (Canonical, London, UK) and was equipped with an Intel I9-13900K processor, Intel X520-DA2 network adapter (Intel, Santa Clara, CA), where camera data was saved to a 2 TB Samsung 990 SSD (Samsung, Suwon-si, South Korea). The desktop machine synchronized all clocks in the network as PTP master, controlled all cameras, stored down acquired data from each camera, and acquired neural data over Bluetooth from a proprietary Neuralink SDK.

To accommodate the high data rates associated with streaming high-resolution frames from multiple devices at such high frequencies, all frames were stored in an HDF5 file format with lossless compression filters enabled. Additionally, each camera’s stream was double-buffered to prevent potential collisions in buffer usage and read/write operations.

### Neural Data Processing - Head-fixed Utah Array

Neuronal data from primary motor cortex were recorded using a 96-channel Utah array (Blackrock Microsystems, Salt Lake City, Utah, USA) and acquired through a Blackrock CerePlex E headstage. Raw data were recorded at 30kHz and high-pass filtered using a fourth order, zero phase Butterworth filter with a 250Hz cutoff frequency. Rolling root mean square (RMS) voltages were first calculated for all channels high-passed filtered data, and periods where the RMS values exceeded 40µV were rejected. A baseline RMS value was determined by calculating the average RMS voltage for the first minute of recording. Action potentials were then determined by from threshold crossings that occurred at a multiple of the RMS value; this was -3.5× for both Monkey O and Monkey U. For each identified action potential, segments extending 0.53ms before and 1.07ms after each threshold crossing were taken to produce the action potential waveform. These waveforms were then checked to ensure that they possessed physiological characteristics of fast- and regular-spiking cortical neurons. Waveforms with peak-to-trough amplitudes exceeding 300µV were excluded, as were waveforms extending longer than 1ms, which was determined by the time difference between the prominent peaks in the waveform’s temporal derivative. Each channel’s waveforms were pooled, and principal component analysis was used to identify potential artifacts in the recording. Binary spike rasters were determined by the threshold crossing times of the remaining, valid waveforms. These rasters were downsampled to 1kHz and aligned to behavioral data using the LiCoRICE software platform [39].

### Neural Data Processing - Unconstrained Utah Array

From the same Utah array described above, in the unconstrained rig, neuronal data were wirelessly transmitted at 30 kHz using a Cereplex W headstage (Blackrock Microsystems, Salt Lake City, Utah, USA). Raw data were filtered at 250Hz using the same Butterworth filter described above. For all channels, periods where the rolling RMS voltage exceeded 70µV were excluded from analysis. Given the increased presence of artifact in the unconstrained rig, RMS values were calculated by sampling 500, 600ms windows and taking the median RMS value. As with the head-fixed data, actions potentials were determined by threshold crossings at a multiple of the median RMS value (-3.5× for Monkeys O and U) and action potentials with peak-to-trough amplitudes exceeding 300µV or widths greater than 1ms were rejected. Due to the increased presence of small amplitude artifacts in the unconstrained data that were not reliably rejected by PCA filtering, a cross correlation was performed on action potential waveforms against template waveforms collected in the head-fixed rig. This correlation ensured that action potential has a physiologically appropriate shape and that the depolarization period lasted less than 200 ms. As a final check to protect against array-wide artifacts, periods where spiking activity was simultaneously detected on 30 or more channels were rejected.

### Neural Data Processing - N1

For the N1 device, all filtering and spiking event detection was performed on chip using a proprietary algorithm. Spiking events were detected on a per-channel basis, and counts binned at 15 ms were transmitted from the device via Bluetooth. These events were timestamped using PTP, allowing them to be easily synchronized with depth camera recordings.

### Low-dimensional Projections

Spiking data from all identified neural epochs were smoothed using a Gaussian filter with a 30 ms kernel. To generate the shared neural state space, neural epochs were combined across tasks. Standard PCA was performed on the combined data to reduce dimensionality, with equal amounts of data taken from each task. Neural trajectories were then projected onto the desired number of PC dimensions.

### Context angle analysis

To find the angle separating low-dimensional activity between tasks (or between tasks and PC 1), task activity was first projected using PCA as described above. PCA was again performed on each group of task activity, with the first PC providing an N-dimensional vector along the main axis of variance. The angle was calculated by taking the the dot product of the desired vectors using the formula

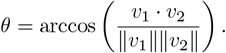

This angle was then converted from radians to degrees.

### Context overlap

To quantify the similarity between contexts, the data were restricted to the desired number of dimensions (2-6). The Mahalanobis distance was calculated for all points for each context, and points with distances greater than the 95th percentile were excluded from analysis. From the remaining points, the N-dimensional convex hulls were created using the scipy Python package (v1.15.3) [65]. The spatial overlap between the two convex hulls (*H*_1_ and *H*_2_) was quantified by calculating their volumetric Jaccard index, referred to as the overlap ratio. This ratio is defined as the volume of the intersection of the two hulls divided by the volume of their union.

The calculation is expressed by the formula:

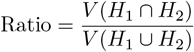

where *V* (*H*_1_ ∩ *H*_2_) is the volume of the intersection and *V* (*H*_1_ ∪ *H*_2_) is the volume of the union. The volume of the union was calculated using the principle of inclusion-exclusion: *V* (*H*_1_ ∪ *H*_2_) = *V* (*H*_1_) + *V* (*H*_2_) − *V* (*H*_1_ ∩ *H*_2_).

Substituting this into the primary equation yields the final formula used for the calculation:

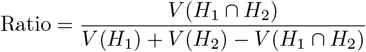

Here, *V* (*H*_1_) and *V* (*H*_2_) are the volumes of the individual convex hulls, and the intersection volume *V* (*H*_1_ ∩ *H*_2_) was approximated using a Monte Carlo sampling method.

### Context shift quantification

To quantify the shift as Monkeys H and P switched between contexts, the PCA projected neural activity from the mouse-reaching and free movement tasks was taken and used to calculate the 20-dimensional centroids for these tasks. Preceding and following the mouse-reaching and computer tasks, animals were in the section of their enclosure that the free movement task occurred. Neural activity 60 seconds preceding and 60 seconds following the start or completion of the mouse-reaching task was examined. This insured that periods of seated reaching and free movement were captured. For all timepoints of neural activity that fell within this window, a difference metric was computed as follows:

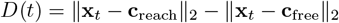

where *D*(*t*) is the difference metric at a given timepoint *t*; x_*t*_ is the 20-dimensional PCA-projected neural activity vector at time *t*; c_reach_ and c_free_ are the 20-dimensional centroids for the reaching and free tasks, respectively; and ∥ · ∥_2_ denotes the L2 norm. For this difference metric, positive values indicate activity closer to the free task centroid, while negative values indicate activity closer to the reaching task centroid.

### PCA dimensionality

PCA dimensionality was calculated as the fewest PCs needed to capture ≥ 80% of the total variance, unless specified otherwise. To characterize how dimensionality scales with the number of channels, analyses on channel subsets were performed. Channel counts were sampled from 10 to 90 for Monkeys O and U, and from 20 to 1000 (in increments of 10) for Monkeys H and P. For each recording session and task, subsampling was performed 1000 times with replacement for each count. Linear, logarithmic, and power law models were evaluated to describe this relationship. Power law and linear fits best modeled the data, and model fits across animals can be found in Table S2.

### Factor analysis dimensionality

Neural dimensionality was estimated using Factor Analysis (FA) implemented in the Python scikit-learn package [45]. Neural data were preprocessed as previously described, and all channels were standardized using a z-score transformation.

The optimal number of latent factors was determined by minimizing the Bayesian Information Criterion (BIC). For a given dataset for *N* subsampled channels, independent FA models were fit across a range of latent dimensions *d*. For Monkeys O and U, channel sets from 20 to 90 (incrementing by 10) were swept, and latent dimensions from 2 to *N/*2 (incrementing by 2) were tested. For Monkeys H and P, channel sets from 100 to 1000 (incrementing by 50) were swept, and latent dimensions from 10 to *N/*2 (incrementing by 10) were tested. Model parameters were estimated using randomized Singular Value Decomposition (SVD). For each model, the BIC was computed as:

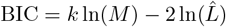

where 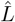 is the maximum log-likelihood of the data and *k* represents the number of free parameters in the Factor Analysis model, calculated as:

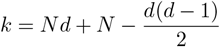

The intrinsic dimensionality was identified as the value of *d* that minimized the BIC. This procedure was bootstrapped over 100 iterations for each value of *N* channels.

### Shared variance component analysis

Shared variance component analysis (SVCA), is described in detail by Stringer et al. [63], and SVCA calculations were performed using a Python implementation (https://github.com/JoramKeijser/SVCA). In brief, neural activity from the free-behavior task was binned, smoothed and z-scored. To perform SVCA, channels were split into two non-overlapping subsets (*F* and *G*). The data were further split into training and testing sets using interleaved 15-second blocks to cross-validate covariance over time. Maximum covariance analysis was computed on the training data by computing the singular value decomposition of the cross-covariance matrix between the two subsets 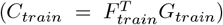. This yielded pairs of projection axes (shared variance components) that maximized the covariance between the two groups.

The reliability of these components was quantified by projecting the held-out testing data onto these training-derived axes and computing the resulting covariance. To determine significant dimensionality, we compared this variance against a null distribution generated by shuffling the temporal alignment of the neuron subsets. Adapting the approach used by Manley et al. [37], a component was deemed significant if its variance exceeded the mean of the shuffled distribution by at least four standard deviations. The total number of significant components defined the SVCA dimensionality.

For Monkeys O and U, random channel subsets were sampled from 20 to 90 (in increments of 10), while for Monkeys H and P, subsets were sampled from 100 to 1000 (in increments of 50). This procedure was repeated 100 times for each subset size to generate a dimensionality distribution.

### Autoencoder dimensionality estimation

We additionally estimated intrinsic dimensionality using a non-linear autoencoder-based approach. An autoencoder learns to compress high-dimensional input data through a low-dimensional bottleneck and then reconstruct the original input; the reconstruction error at a given bottleneck size reflects how well the data can be represented in that number of dimensions. Sweeping the bottleneck size and identifying the point at which reconstruction error ceases to improve meaningfully produces a non-linear estimate of intrinsic dimensionality.

The autoencoder consisted of a symmetric encoder–decoder architecture with three fully connected layers in each half. For a given channel subset size, the encoder mapped the *N* -dimensional input through hidden layers of size *h*_1_ = max(128, ⌊0.75*N* ⌋) and *h*_2_ = max(64, ⌊0.50*N* ⌋) to a bottleneck layer of size *b*, where both *h*_1_ and *h*_2_ were constrained to be no smaller than *b*. The decoder mirrored this architecture, mapping the bottleneck representation back through layers of size *h*_2_ and *h*_1_ to produce an *N* -dimensional reconstruction. All hidden layers used the Gaussian Error Linear Unit (GELU) activation function; no activation was applied to the bottleneck or output layers.

The input to the autoencoder consisted of spike counts binned at 15 ms and smoothed using a Gaussian filter with a 30 ms kernel. For each analysis, neural data were extracted from 10 non-overlapping temporal windows, centered within the recording session. A window length of 5000 time bins (75 s) was used for Monkey P, while a window length of 3000 time bins (45 s) was used for Monkey H to accommodate the shorter recording duration. For a given channel subset, data within each window were *z*-scored by subtracting the per-channel mean and dividing by the per-channel standard deviation.

Networks were trained for 50 epochs using the Adam optimizer with a learning rate of 10^*−*3^ and weight decay of 10^*−*5^. Mean squared error (MSE) served as the reconstruction loss. Data were split 80/20 into training and validation sets, and training was performed in mini-batches of 256 samples. After training, reconstruction error was evaluated on the held-out validation set.

For each channel subset and temporal window, the bottleneck size *b* was swept from 10 to 300 in increments of 10, excluding values where *b* ≥ *N*. This sweep was repeated across 10 temporal windows, yielding validation MSE curves as a function of bottleneck size. The optimal bottleneck— taken as the non-linear dimensionality estimate—was identified using the Kneedle algorithm, which detects the elbow point on a convex, decreasing curve [53]. This entire procedure was repeated for 100 iterations per channel subset size, with channels randomly sampled without replacement from the full electrode population on each iteration.

### Inverse compression ratio

A detailed explanation of the inverse compression ratio (ICR) can be found in [69]. In brief, ICR is defined as the ratio of the compressed data size to the raw data size:

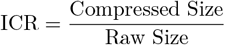

This normalized, unitless metric serves as a proxy for joint entropy, with higher values indicating higher signal complexity (lower regularity). To compress data, the DEFLATE algorithm was utilized (via the standard Python gzip library) on binned, unfiltered spike counts. This method provides a computationally efficient, permutation-sensitive estimate of complexity that can easily be applied to multichannel neural data.

### Multiclass linear discriminant analysis

Multiclass linear discriminant analysis (mcLDA) can be formulated as follows. Let the columns of the data matrix 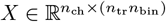 be partitioned into subsets 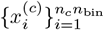 for each class *c* ∈ {1, …, *C*}, where *n*_*c*_ is the number of trials for class *c, n*_bin_ is the number of time bins per trial, and 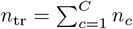 is the total number of trials. Each subset represents neural activity samples recorded during reaches in direction *c*. Define the class-conditional mean as 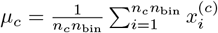. The within-class scatter matrix is then defined as:

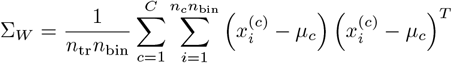

which captures the pooled covariance of the data after removing the effect of class identity. The between-class scatter matrix is defined as:

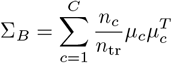

which captures how the class-conditional means are distributed relative to one another. These two matrices satisfy the additive decomposition of total covariance [20]:

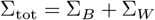

where 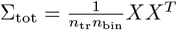 is the total covariance of the (zero-mean) data. The objective of LDA is to find a projection matrix 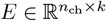 that maps neural population activity to a *k*-dimensional latent space in which classes are maximally separated. This is achieved by maximizing the Fisher criterion:

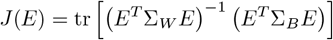

which quantifies the ratio of between-class to within-class variance in the projected space. In practice, ∑_*W*_ may be singular (e.g., if a channel records zero spikes across all trials), preventing direct inversion. To ensure numerical stability, the within-class scatter matrix is regularized as:

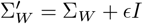

where *ϵ* = 10^*−*5^ is used throughout the analyses presented here. The projection matrix *E* that maximizes *J*(*E*) consists of the *k* eigenvectors corresponding to the largest eigenvalues of 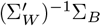. These eigenvectors and their associated eigenvalues are guaranteed to be real-valued, and the rank of the solution space (i.e., the number of positive eigenvalues) is at most *C* − 1, since ∑_*B*_ has rank at most *C* −1. Furthermore, although 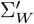 and ∑_*B*_ are individually symmetric, their product 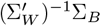 is generally not symmetric, which means the columns of *E* are not inherently orthogonal. To impose orthonormality, Gram–Schmidt orthogonalization is applied to project *E* onto the Stiefel manifold 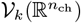 (the set of *n*_ch_ × *k* matrices with orthonormal columns). This does not alter the linear subspace spanned by the columns of *E*, which is the object of primary interest in this framework.

### Trajectory decoding

Neural trajectories were extracted beginning at reach onset. For Monkeys O and U, these raw trajectories spanned 570 ms (38 time bins); for Monkeys H and P, they were truncated to 390 ms (26 time bins) to accommodate the higher variability in reach durations and ensure a consistent number of trials across animals.

Decoding was performed on randomly sampled channel subsets. Subset sizes ranged from 20 to 90 channels in steps of 10 for Monkeys O and U, and from 20 to 1000 channels in steps of 20 for Monkeys H and P. For each subset size, 1000 random channel samples were drawn. Within each sample, the decoded portion of the neural trajectory began 60 ms (4 time bins) after reach onset, and its length was varied systematically. Trajectory length ranged from 15 ms (1 time bin) up to 510 ms (34 time bins) for Monkeys O and U, and up to 330 ms (22 time bins) for Monkeys H and P.

For each channel sample, reach trials were partitioned into training and testing sets using a 90/10 split. A PCA subspace was constructed from the smoothed neural data of the training set. For PCA decoding, a Gaussian Naive Bayes classifier was trained directly on the top 20 PCs, which we found to optimize performance (Figure S7). For mcLDA decoding, the number of retained PCs was capped at 50 for Monkeys O and U and 100 for Monkeys H and P; if the channel subset contained fewer channels than this cap, all PCs were retained. A 7-dimensional mcLDA subspace was then constructed from these PC trajectories and used to train a Gaussian Naive Bayes classifier. In both cases, subspace transforms and classifiers fit on the training set were applied to the test set to compute decoding accuracy.

Performance for each channel-sample and trajectory-length pair was averaged across 10 random train/test splits. This procedure was repeated for all 1000 channel samples at each subset size, and mean accuracy across samples is reported.

### Silhouette score

To evaluate the consistency and separation of the identified clusters, we computed the silhouette score [50]. The silhouette value *s*(*i*) for each data point *i* measures how similar the point is to its own cluster compared to other clusters.

For a given point *i*, let *a*(*i*) be the mean distance between *i* and all other points in the same cluster. Let *b*(*i*) be the minimum mean distance from *i* to all points in any other cluster, defined as:

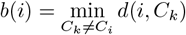

where *d*(*i, C*_*k*_) represents the mean distance between point *i* and all points in cluster *C*_*k*_. The silhouette value for point *i* is then given by:

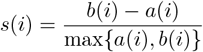

The value of *s*(*i*) ranges from -1 to 1. A value close to 1 implies that the data point is well-matched to its own cluster and poorly matched to neighboring clusters. A value near 0 indicates that the point is on or very close to the decision boundary between two neighboring clusters. While not relevant to our data, a negative value would suggest that the point may have been assigned to the wrong cluster.

Random channel subsets ranging from 50 to 1000 in increments of 10 were used to calculate silhouette scores for all trajectory points. The overall quality of the clustering solution was assessed using the median silhouette score across all samples, providing a global measure of cluster validity.

## Supplementary materials

**Table S1:** Event counts across experimental sessions. These counts represent 500 ms epochs identified for seated reaching and unconstrained movement tasks. Sit, walk, reach, turn, and stand movements all refer to unconstrained behavior.

| Session ID | Seated reach | Sit | Walk | Reach | Turn | Stand |
| --- | --- | --- | --- | --- | --- | --- |
| O240718 | 604 | 38 | 453 | 512 | 449 | - |
| O240719 | 600 | 0 | 401 | 387 | 392 | - |
| O240813 | 574 | 24 | 380 | 332 | 331 | - |
| O240814 | 458 | 10 | 577 | 544 | 561 | - |
| U240610 | 500 | 4 | 101 | 91 | 81 | - |
| U240611 | 277 | 145 | 963 | 892 | 966 | - |
| U240612 | 546 | 107 | 1220 | 1211 | 1035 | - |
| U240613 | 552 | - | - | - | - | - |
| U240614 | 641 | 124 | 1711 | 1524 | 1635 | - |
| H240729 | 189 | 2368 | 151 | - | - | 351 |
| P240608 | 230 | 2592 | 315 | - | - | 1218 |

**Fig. S1:**
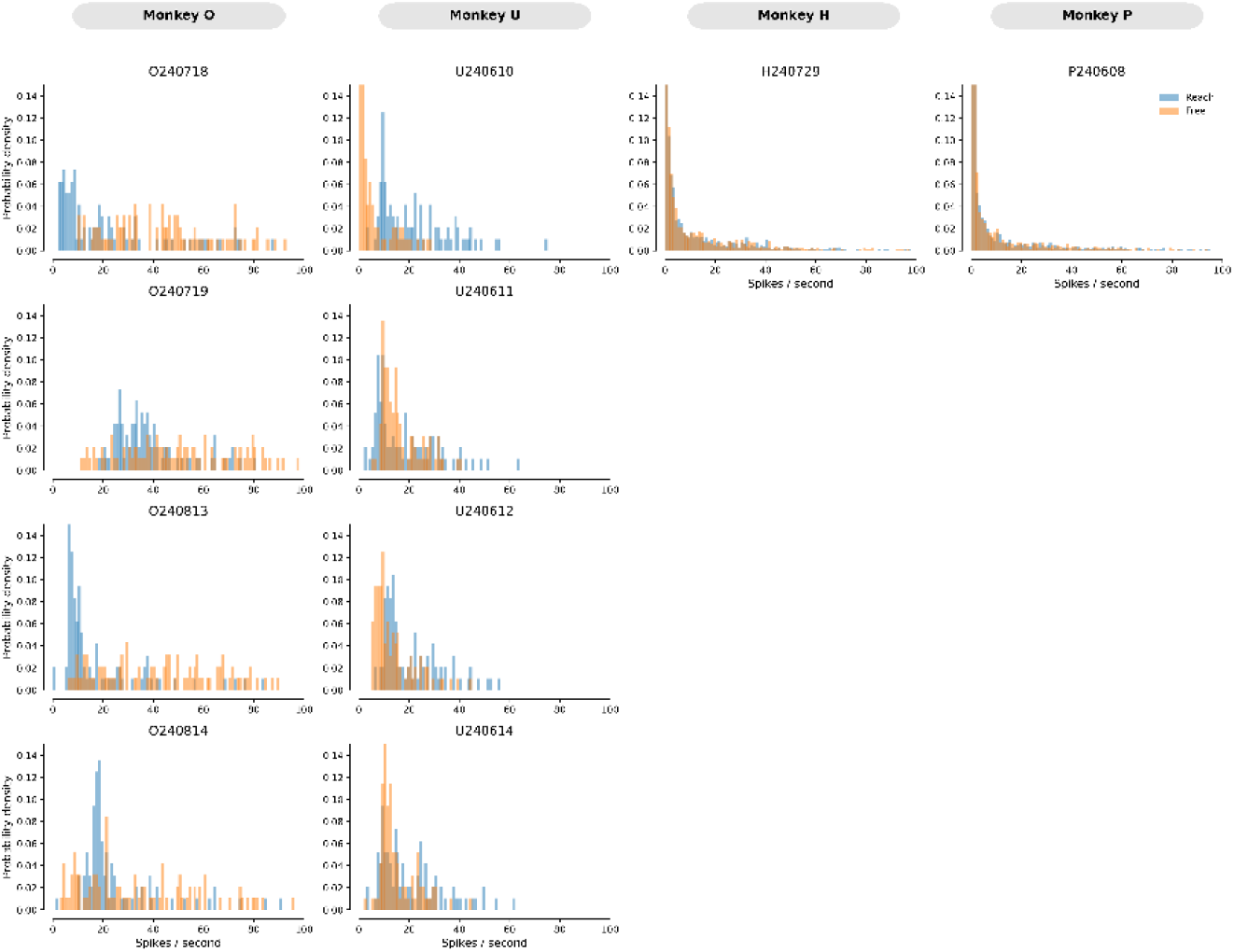
Average firing rates across array channels. For each recording session, firing rates are reported for both seated reaching and free movement tasks. Rates were calculated as the per channel average firing rate across the task session.

**Fig. S2:**
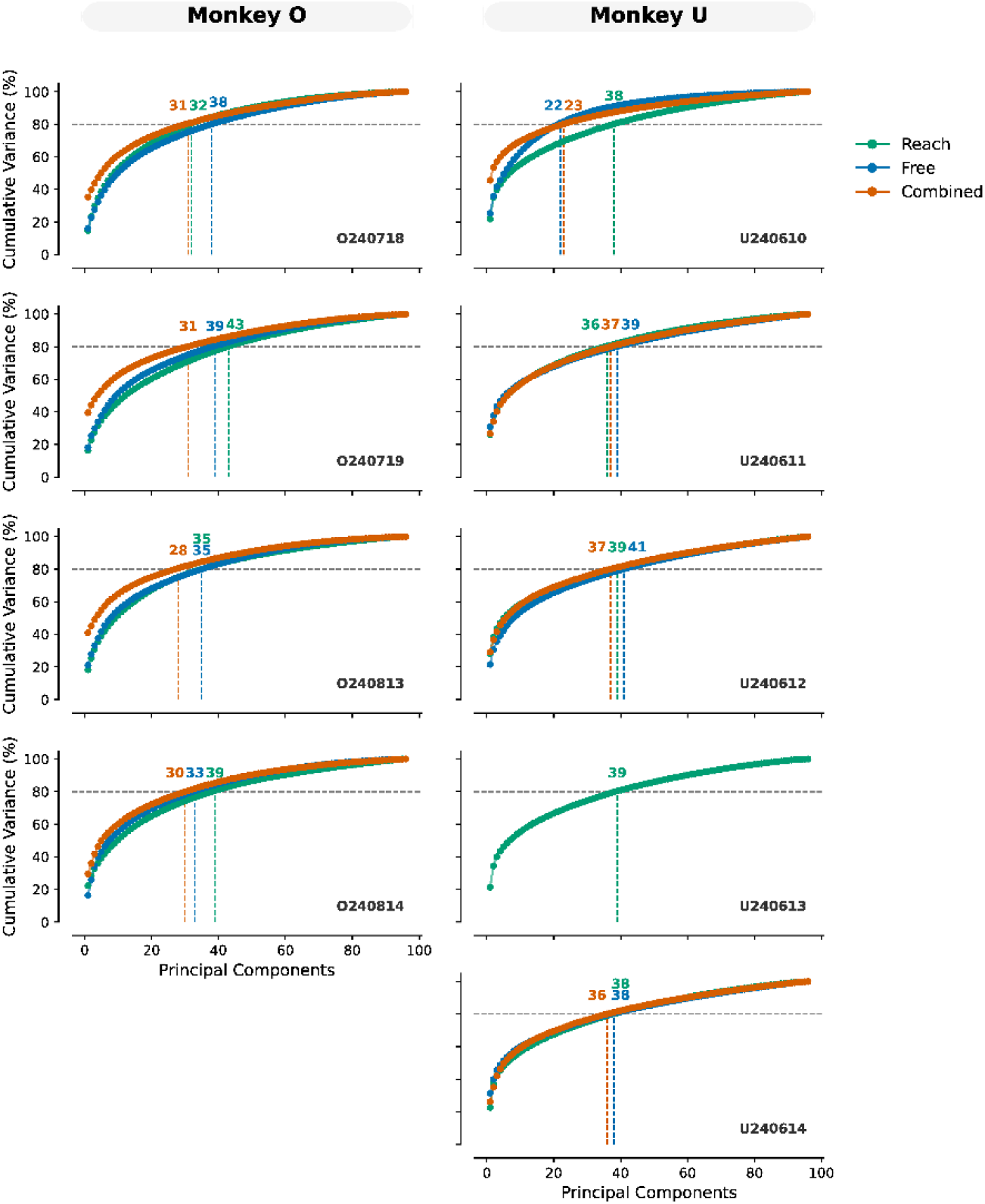
Cumulative variance plots for reaching task (green), freely-moving task (blue), and combined reaching/freely-moving tasks (orange) over individual recording sessions (Monkeys O and U). Recording date (prefixed with the first letter of the animal’s name) is located in the bottom right of each subplot. The number of dimensions needed to achieve 80 percent of the total variance is shown for each trace.

**Fig. S3:**
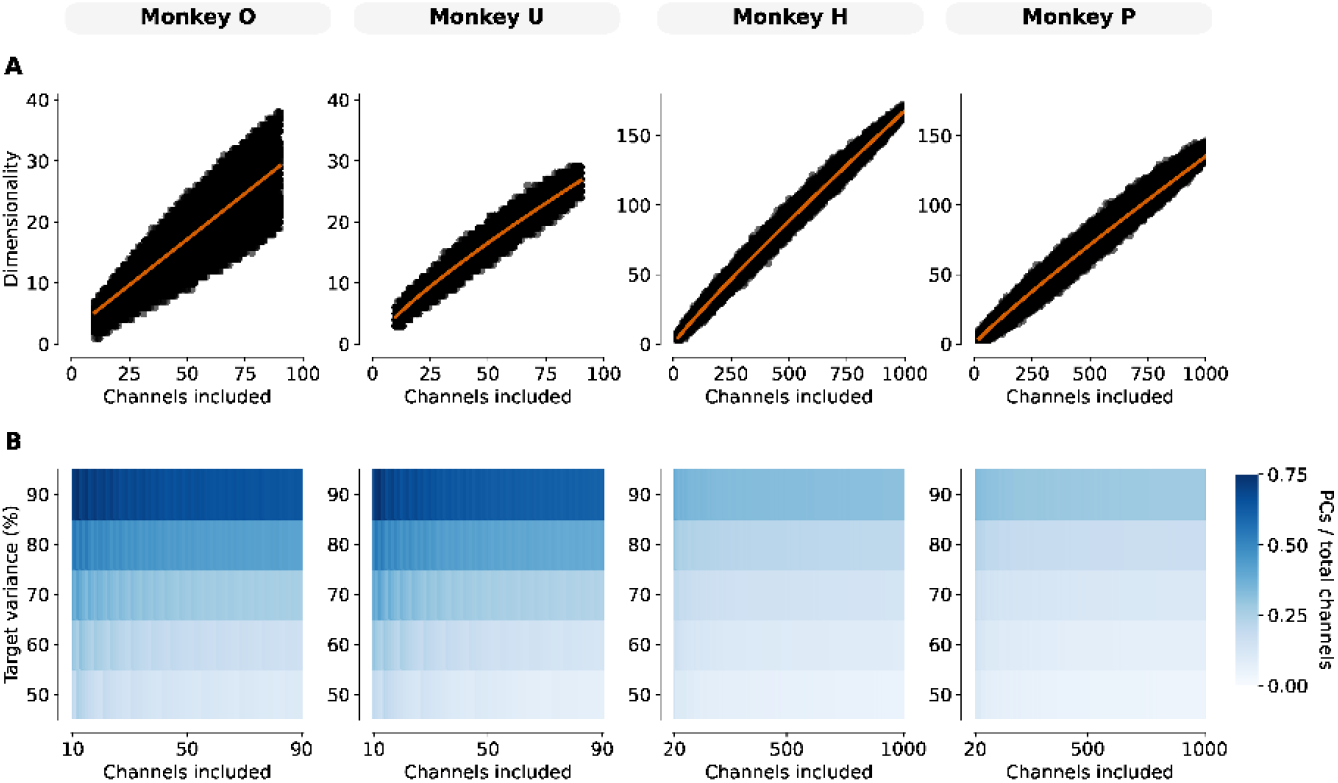
Variance plots for scated reaching task. (**A**) and (**B**) mirror Figure 2 (**B**) and (**C**) but are calculated on scated reaching task data. Fit lines represent the linear or power law model that best described the data (see Table S2).

**Table S2:**
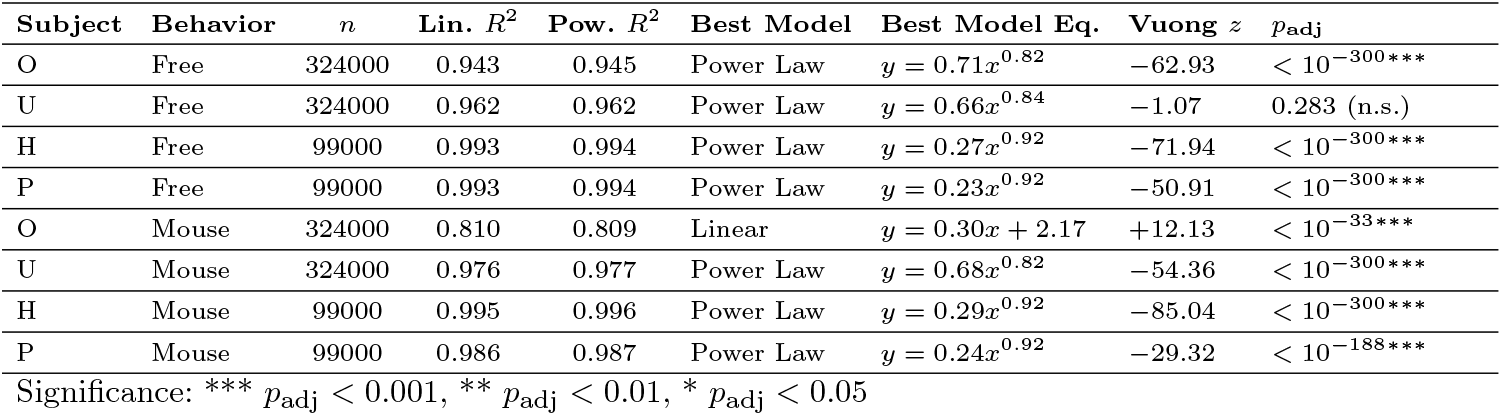
Comparison of linear vs. power-law fits for the scaling of unsupervised dimensionality with electrode count. The number of bootstrap dimensionality estimates is given by *n*, and the residual degrees of freedom for all calculations is *n* − 2. Best model is selected by a two-sided Vuong closeness test; positive *z* favors the linear model, negative *z* favors the power law. Reported *p* values are Benjamini–Hochberg FDR-adjusted across the eight tests (animal × behavior). 95% CIs for all regression coefficients are provided in Table S3.

| Subject | Behavior | $n$ | Lin. $R^2$ | Pow. $R^2$ | Best Model | Best Model Eq. | Vuong $z$ | $p_{\text{adj}}$ |
| --- | --- | --- | --- | --- | --- | --- | --- | --- |
| O | Free | 324000 | 0.943 | 0.945 | Power Law | $y = 0.71x^{0.82}$ | -62.93 | $< 10^{-300***}$ |
| U | Free | 324000 | 0.962 | 0.962 | Power Law | $y = 0.66x^{0.84}$ | -1.07 | 0.283 (n.s.) |
| H | Free | 99000 | 0.993 | 0.994 | Power Law | $y = 0.27x^{0.92}$ | -71.94 | $< 10^{-300***}$ |
| P | Free | 99000 | 0.993 | 0.994 | Power Law | $y = 0.23x^{0.92}$ | -50.91 | $< 10^{-300***}$ |
| O | Mouse | 324000 | 0.810 | 0.809 | Linear | $y = 0.30x + 2.17$ | +12.13 | $< 10^{-33***}$ |
| U | Mouse | 324000 | 0.976 | 0.977 | Power Law | $y = 0.68x^{0.82}$ | -54.36 | $< 10^{-300***}$ |
| H | Mouse | 99000 | 0.995 | 0.996 | Power Law | $y = 0.29x^{0.92}$ | -85.04 | $< 10^{-300***}$ |
| P | Mouse | 99000 | 0.986 | 0.987 | Power Law | $y = 0.24x^{0.92}$ | -29.32 | $< 10^{-188***}$ |

Significance: \*\*\* $p_{\text{adj}} < 0.001$ , \*\* $p_{\text{adj}} < 0.01$ , \* $p_{\text{adj}} < 0.05$

**Table S3:** Statistical details for the best-fitting dimensionality scaling models. Equations include 95% confidence intervals for each model equation coefficient.

| Subject | Behavior | Best Model | Best Model Eq. [95% CI] |
| --- | --- | --- | --- |
| O | Free | Power Law | $y = (0.71 [0.71, 0.71])x^{(0.82 [0.81, 0.82])}$ |
| U | Free | Power Law | $y = (0.66 [0.66, 0.66])x^{(0.84 [0.84, 0.84])}$ |
| H | Free | Power Law | $y = (0.27 [0.27, 0.27])x^{(0.92 [0.92, 0.92])}$ |
| P | Free | Power Law | $y = (0.23 [0.23, 0.23])x^{(0.92 [0.92, 0.92])}$ |
| O | Mouse | Linear | $y = (0.30 [0.30, 0.30])x + (2.17 [2.14, 2.20])$ |
| U | Mouse | Power Law | $y = (0.68 [0.68, 0.68])x^{(0.82 [0.82, 0.82])}$ |
| H | Mouse | Power Law | $y = (0.29 [0.29, 0.29])x^{(0.92 [0.92, 0.92])}$ |
| P | Mouse | Power Law | $y = (0.24 [0.24, 0.24])x^{(0.92 [0.92, 0.92])}$ |

**Fig. S4:**
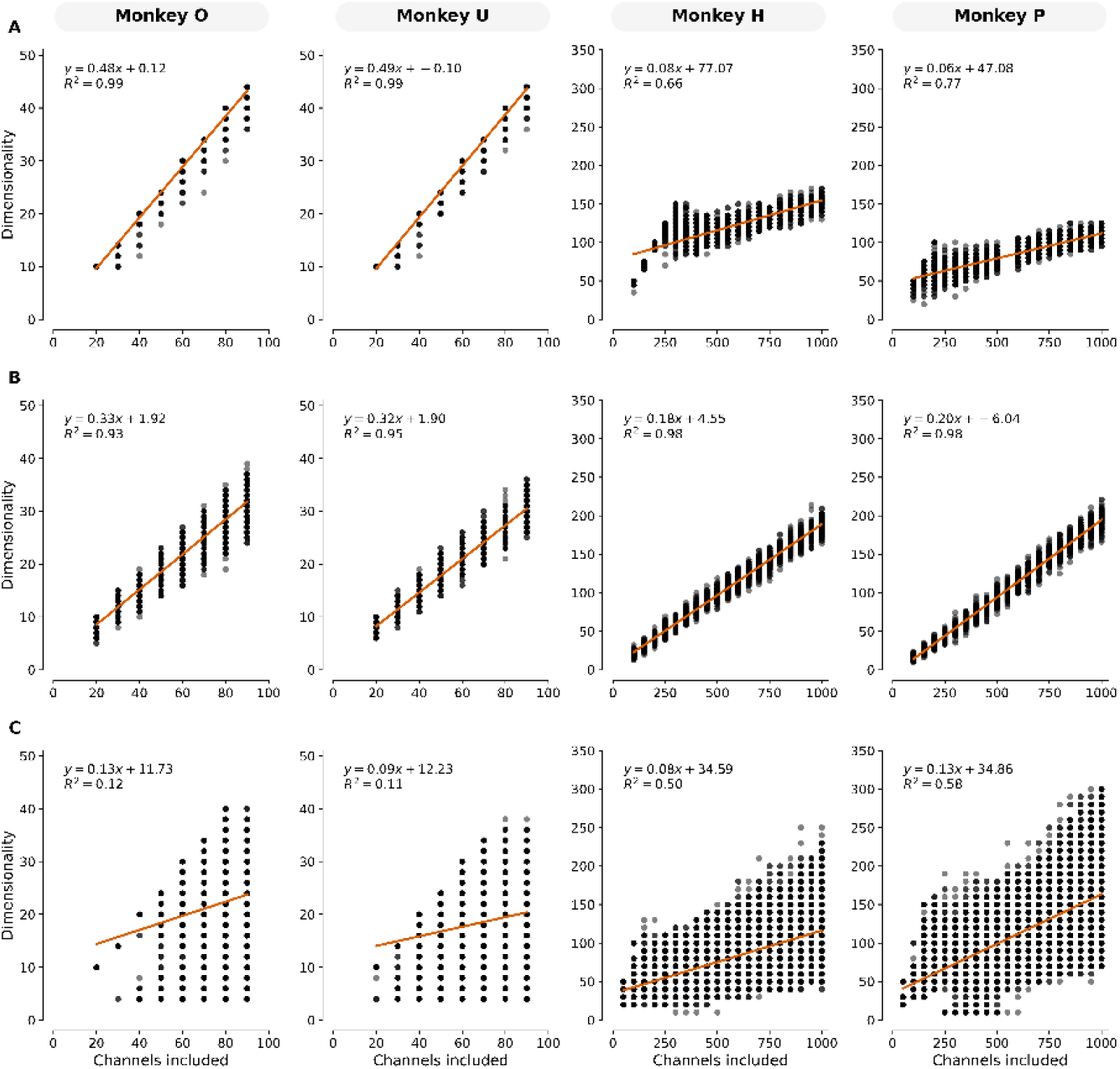
Unsupervised dimensionality metrics scale with channel count. (**A**) Over channel subsets, independent factor analysis models were fit across a range of latent dimensions *d*. The intrinsic dimensionality was identified as the value of *d* that minimized the Bayesian Information Criterion score (sec Methods). Linear fits (orange line) are provided for each animal, along with fit equations and R2 values. (**B**) Same as (**A**), but using shared variance component analysis. Here, variance components were considered significant if they were more than 4 standard deviations above the shuffled distribution. (**C**) Same as (**A**), but using autoencoder models. Here, the autoencoder bottleneck size was used as a proxy for dimensionality. For a particular channel subset on a given window of data, the optimal bottleneck size represents the elbow point of the validation mean squared error as determined by the Kneedle algorithm [53].

**Fig. S5:**
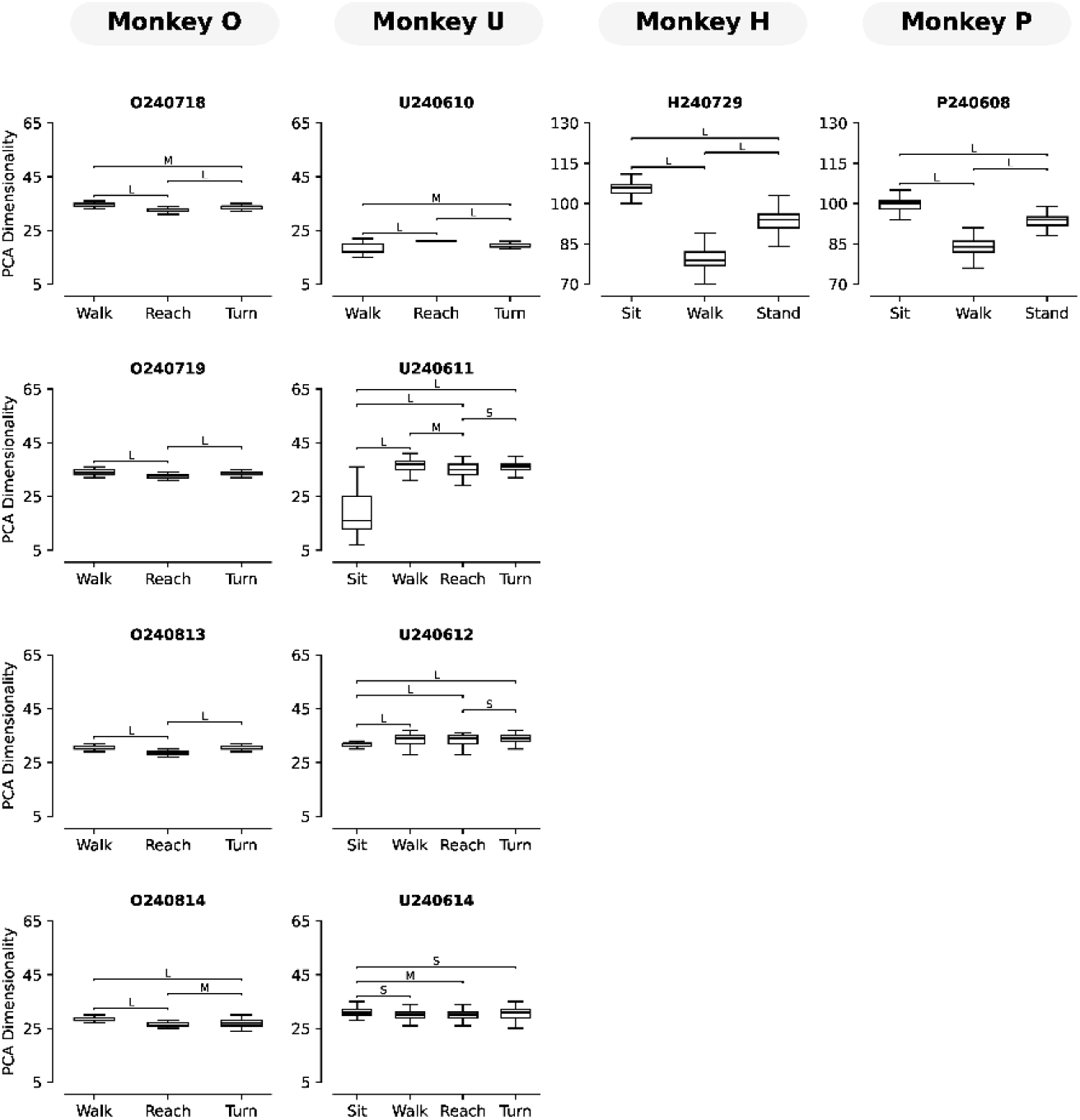
Differentiation of behaviors during free movement tasks using PCA. For each behavior on a given recording session, PCA dimensionalities were calculated on 30-second concatenated windows of raw spiking data. Cliff’s delta was calculated for each within-session pairwise movement comparison, and magnitude thresholds are reported as follows: |*δ*| < 0.147 negligible, 0.147 ≤ |*δ*| < 0.330 small, 0.330 ≤ |*δ*| < 0.474 medium, and |*δ*| ≥ 0.474 large [49]. Across animals, most behavior pairs were distinguished by medium-to-large effect sizes.

**Fig. S6:**
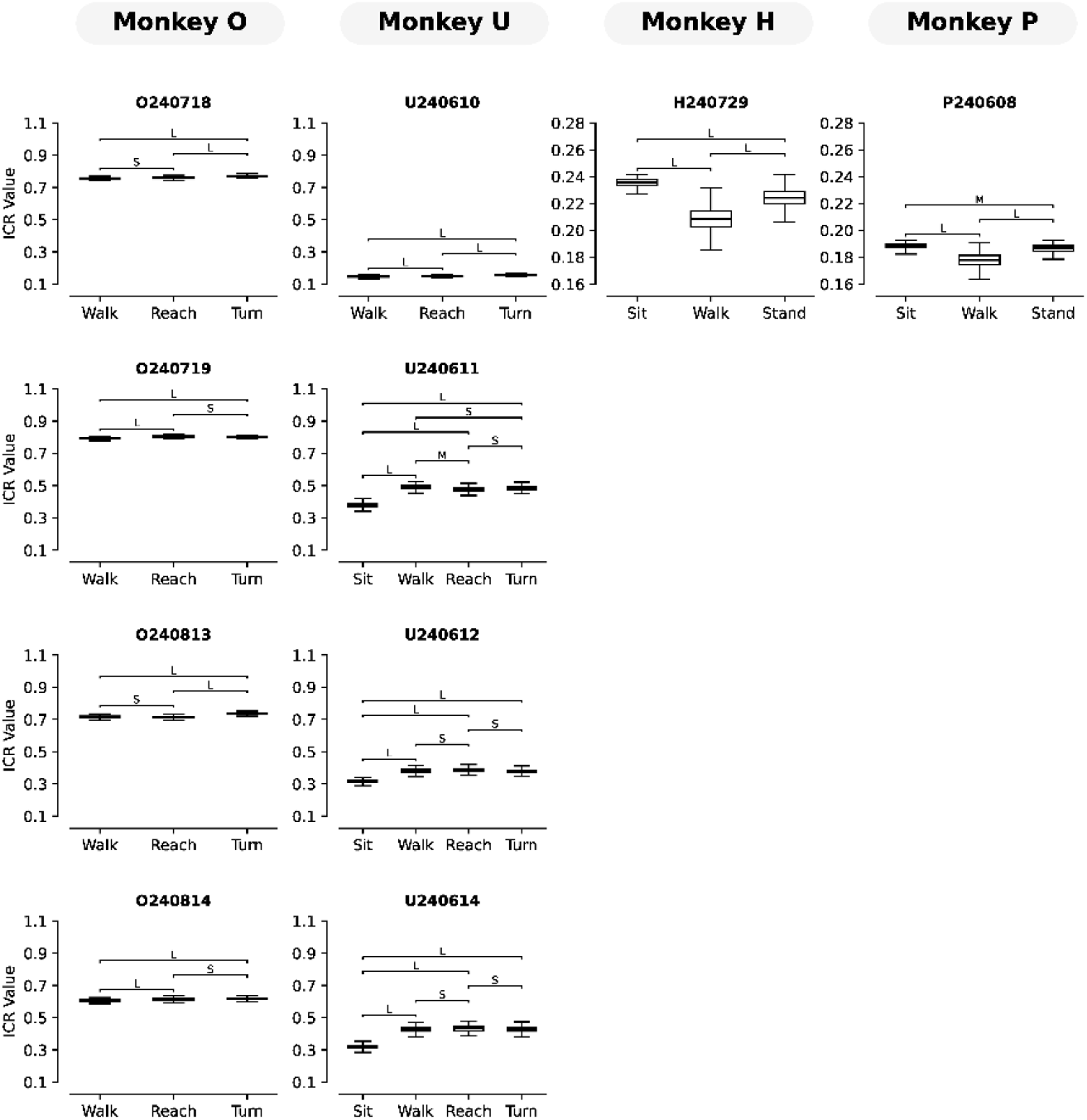
Differentiation of behaviors during free movement tasks using ICR. For each behavior, ICR dimensionalities were calculated on the same 30-second concatenated windows of raw spiking data used in Figure S5. Cliff’s delta was calculated for each within-session pairwise movement comparison, and magnitude thresholds are reported as follows: |*δ*| < 0.147 negligible, 0.147 ≤ |*δ*| < 0.330 small, 0.330 ≤ |*δ*| < 0.474 medium, and |*δ*| ≥ 0.474 large [49]. Compared to PCA, ICR yielded fewer negligible effects across sessions, returning non-negligible effects for nearly all movement pairs.

**Fig. S7:**
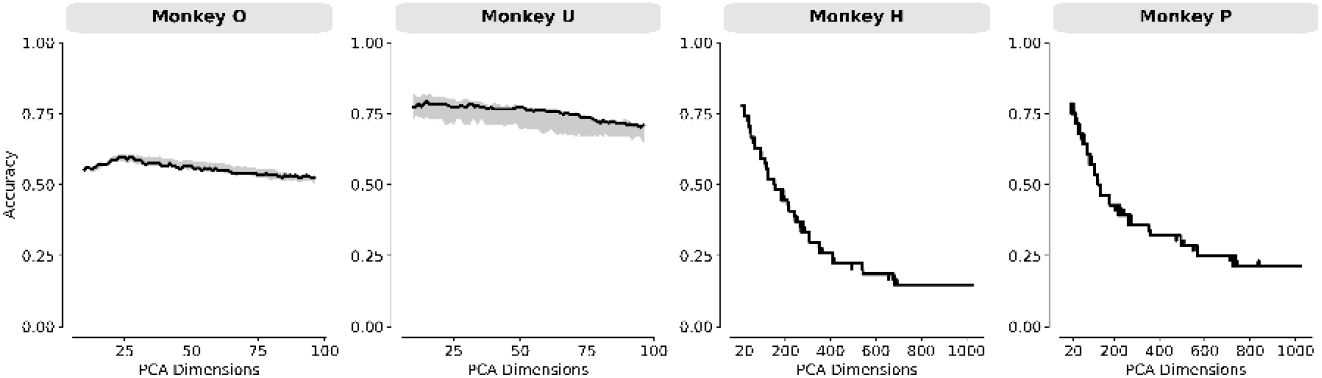
Decoder accuracy as a function of PCA dimensions incorporated. Shading indicates interquartile range. Above 20 PCs, decoding performance drops for all animals.

**Fig. S8:**
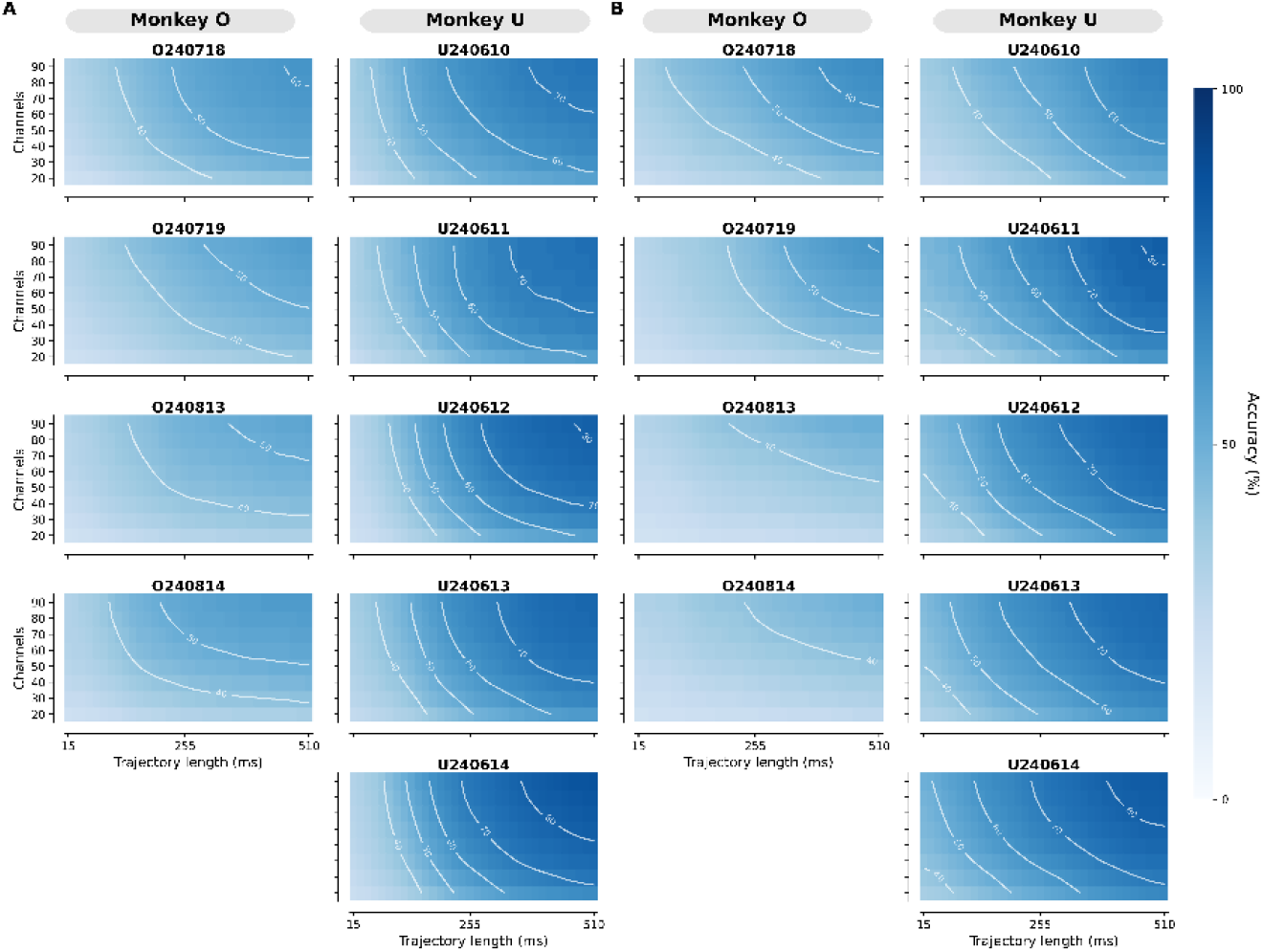
Decoding performance on individual recording sessions for Monkeys O and U. Recording date (prefixed with the first letter of the animal’s name) is located above each plot. (**A**) Decoding performance using PCA trajectories. (**B**) Decoding performance using mcLDA trajectories.

**Fig. S9:**
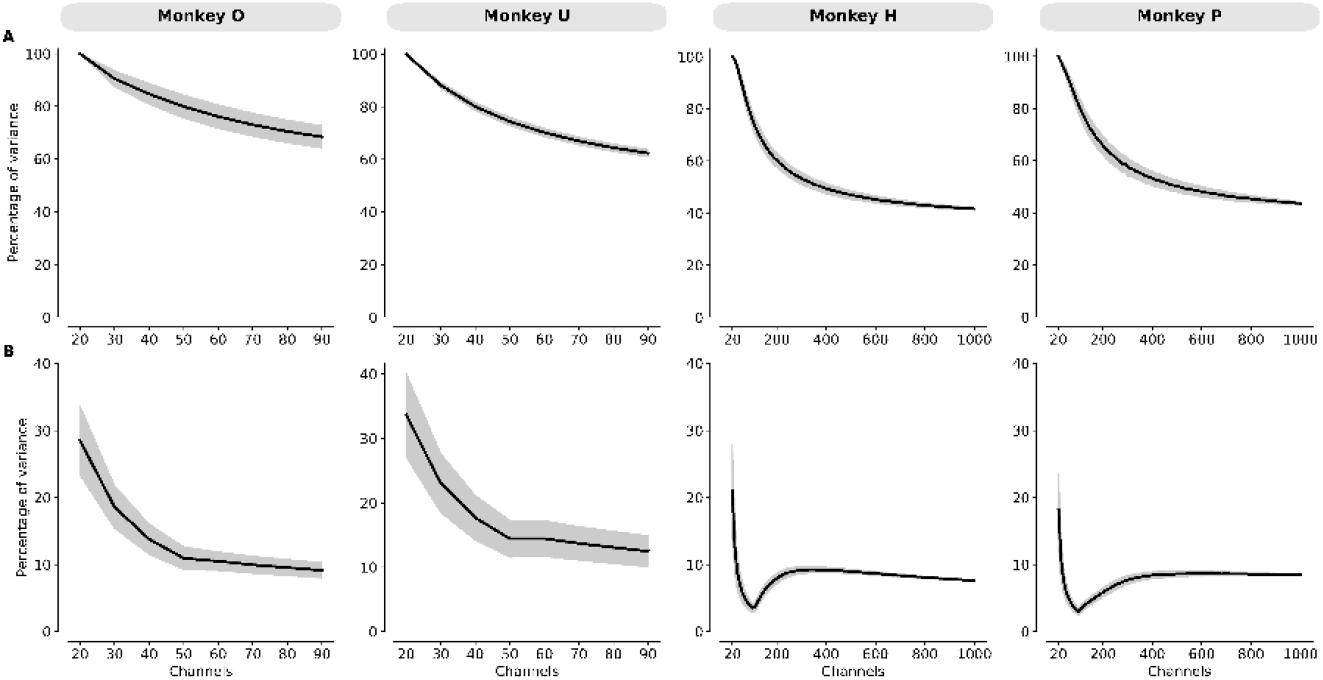
Percentage of total variance captured by PCA and mcLDA models. (**A**) Percentage of total variance captured by the top 20 PCs used for PCA decoding. Lines show means across 1000 channel subsamples per recording session; shading indicates ±1 standard deviation. Captured variance decreases as more channels are included, but even at the maximum channel counts, these PCs still explain over 40% of the total variance. (**B**) Percentage of total variance captured by the 7 mcLDA dimensions, computed as the product of two factors: the variance captured by the input PC features and the fraction of that PC subspace reflected in the mcLDA dimensions. Sampling and shading conventions as in (**A**). The mumber of PC features supplied to mcLDA was capped at 50 for Monkeys O and U and 100 for Monkeys Hand P. For all animals, captured variance drops sharply as PC features are added up to this cap. Beyond the cap, Monkeys and U show a gradual decline with additional channels, while Monkeys H and I show a brief rebound–reflecting mcLDA extracting more discriminative information from the fixed PC pool as channel count grows–followed by a gradual decline. At the maximum channel counts, incLDA uses 5-6× less variance than PCA.

**Fig. S10:**
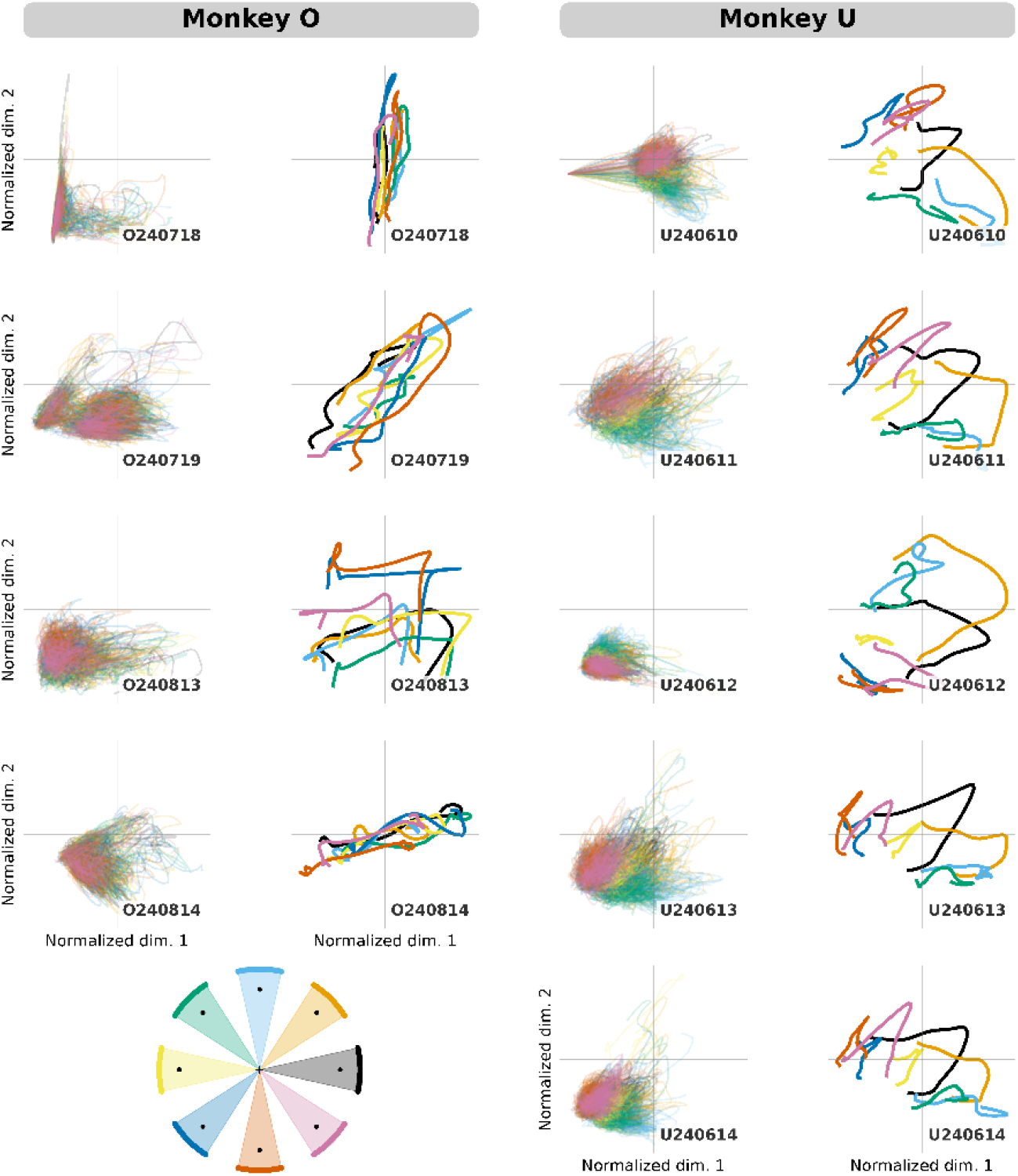
Individual and averaged radial-8 PCA reaching trajectories for Monkeys O and U. For each monkey, individual PCA trajectories (left column) and trajectories averaged by direction (right column) are shown. Trajectory colors correspond to reach direction, as shown in the graphic (bottom left). Neither individual nor averaged PCA trajectories are well separated by direction.

**Fig. S11:**
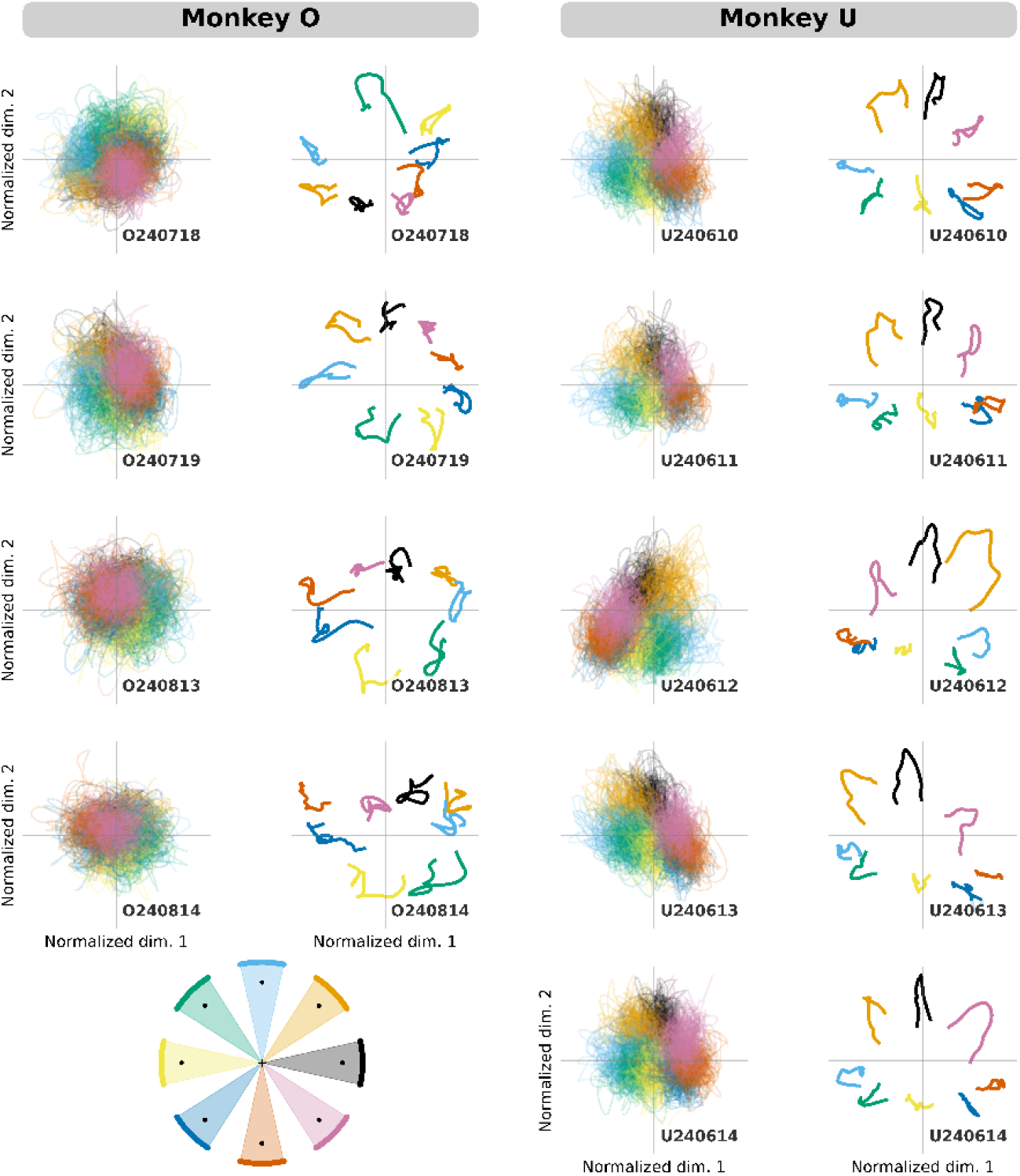
Individual and averaged radial-8 mcLDA reaching trajectories for Monkeys O and U. For cach monkey, individual meLDA trajectories (left column) and trajectories averaged by direction (right column) are shown. Trajectory colors correspond to reach direction, as shown in the graphic (bottom left). While individual trajectories are not well separated by direction, this separation appears when trajectories are averaged.

**Fig. S12:**
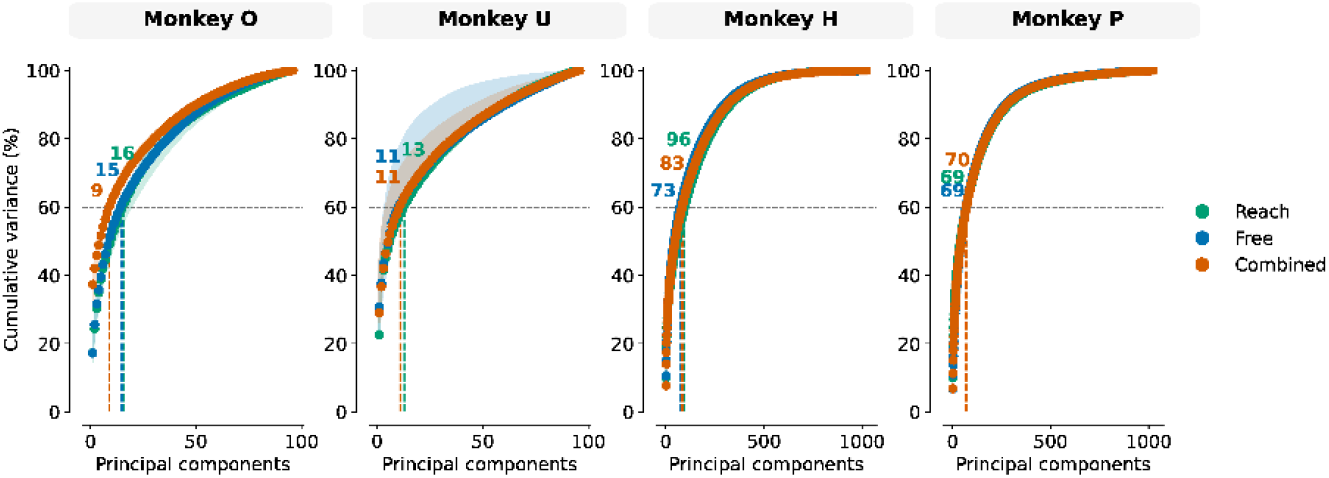
Dimensionality calculations using a 60 percent threshold, as used in [22]. This produces lower dimensionalities which align with previously reported estimates.

**Fig. S13:**
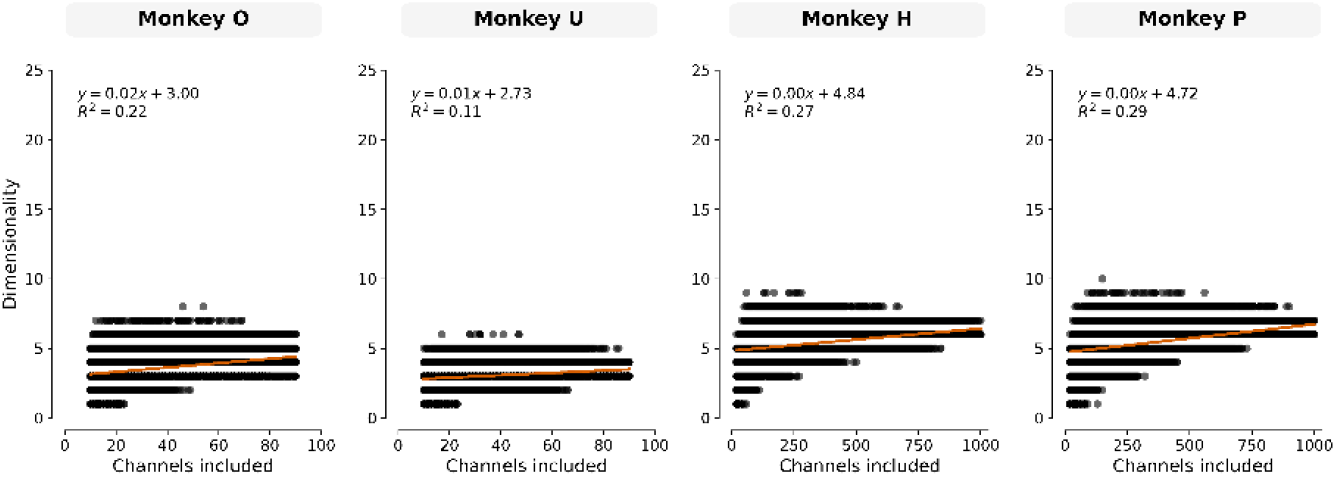
PCA dimensionality when using trial-averaged trajectories. Compared to single-trial analysis, trial-averaged dimensionalities are substantially lower. Scaling is still present as channel count increases, with linear models providing the best fit.

